# Rapamycin exerts its geroprotective effects in the ageing human immune system by enhancing resilience against DNA damage

**DOI:** 10.1101/2025.08.15.670559

**Authors:** Loren Kell, Eleanor J Jones, Nima Gharahdaghi, Daniel J Wilkinson, Kenneth Smith, Philip J Atherton, Anna K Simon, Lynne S Cox, Ghada Alsaleh

## Abstract

mTOR inhibitors such as rapamycin are among the most robust life-extending interventions known, yet the mechanisms underlying their geroprotective effects in humans remain incompletely understood. At non-immunosuppressive doses, these drugs are senomorphic, i.e. they mitigate cellular senescence, but whether they protect genome stability itself has been unclear. Given that DNA damage is a major driver of immune ageing, and immune decline accelerates whole-organism ageing, we tested whether mTOR inhibition enhances genome stability. In human T cells exposed to acute genotoxic stress, we found that rapamycin and other mTOR inhibitors suppressed senescence not by slowing protein synthesis, halting cell division, or stimulating autophagy, but by directly reducing DNA lesional burden and improving cell survival. *Ex-vivo* analysis of aged immune cells from healthy donors revealed a stark enrichment of markers for DNA damage, senescence, and mTORC hyperactivation, suggesting that human immune ageing may be amenable to intervention by low-dose mTOR inhibition. To test this *in vivo*, we conducted a placebo-controlled experimental medicine trial in older adults administered with low-dose rapamycin. p21, a marker of DNA damage-induced senescence, was significantly reduced in immune cells from the rapamycin compared to placebo group. These findings reveal a previously unrecognised role for mTOR inhibition: direct genoprotection. This mechanism may help explain rapamycin’s exceptional geroprotective profile and opens new avenues for its use in contexts where genome instability drives pathology, ranging from healthy ageing, clinical radiation exposure, and even the hazards of cosmic radiation in space travel.

## INTRODUCTION

Rapamycin and other mTOR inhibitors used at low doses increase lifespan in all species tested to date (Bjedov et al., 2010, Ha and Huh, 2011, Harrison et al., 2009). Importantly, this lifespan extension corresponds with increased healthspan, as rapamycin has been shown to improve health across multiple domains (Wilkinson et al., 2012). Further gains in lifespan extension have been reported when rapamycin is administered in combination with other geroprotectors such as trametinib (Gkioni et al., 2025).Though mTOR inhibitors have shown remarkable anti-ageing potential, the exact hallmarks of ageing on which they impact are not fully understood (Weichhart, 2018). One explanation is that mTOR inhibitors such as rapamycin are senomorphic, in that they limit cellular senescence, a physiological process by which highly damaged cells exit the cell cycle and assume a pro-inflammatory, tissue-remodelling phenotype (Walters et al., 2016, Rolt et al., 2019, Park et al., 2020, Walters and Cox, 2018). mTOR activity increases during the *in vitro* senescence of primary human fibroblasts and in human muscle ageing *in vivo* ((Walters et al., 2016, Carroll et al., 2017, Markofski et al., 2015). Further to this correlative data, cells with constitutive mTOR activation enter premature replicative cell senescence *in vitro*, suggesting mTOR hyperactivity is sufficient to drive cellular ageing (Zhang et al., 2003). Consistent with a role of mTOR in ageing and senescence, mTOR inhibitors attenuate a variety of senescence phenotypes and extend replicative lifespan *in vitro* (Walters et al., 2016, Rolt et al., 2019, Park et al., 2020). In humans, rapamycin reduced the presence of dermal cells expressing the senescence biomarker, p16, when administered in a topical skin cream (Chung et al., 2019). The primary cellular mechanism underlying these senomorphic properties of mTOR inhibition are not fully understood, though impacts on slowing protein synthesis, the cell cycle, or supporting the removal of dysfunctional organelles and protein aggregates through enhanced autophagy have been suggested (Weichhart, 2018). Furthermore, there is a gap in our understanding of how mTOR activity is associated with the ageing of cells which drive the ageing process – namely, those of the immune system, for which there is increasing evidence that DNA damage is a key driver (Kell et al., 2023).

Recent studies have demonstrated how ageing of the immune system (immunosenescence) can precipitate whole-organism ageing (Yousefzadeh et al., 2021b, Desdin-Mico et al., 2020), highlighting how strategies which target immunosenescence are at the frontiers of geriatric medicine. Since aged T cells drive tissue destruction and multimorbidity during ageing, they further provide a cellular target for therapeutic anti-ageing intervention (Soto-Heredero et al., 2023). At high doses, rapamycin is immunosuppressant and causes side effects such as poor wound healing, ulcers, and loss of metabolic control leading to diabetes (Knight et al., 2007, Altomare et al., 2006, Houde et al., 2010). On the other hand, at low doses, mTOR inhibition is one of the few interventions which has been shown actually to improve immunity in older people – i.e., to attenuate immunosenescence (Mannick and Lamming, 2023). In humans, low-dose mTOR inhibitor RAD001 (everolimus) improved B and T cell responses to influenza vaccination in older adults (Mannick et al., 2018, Mannick et al., 2014). A second generation mTOR inhibitor RTB101 significantly reduced respiratory tract infections (RTIs) in older adults in a Phase 2b clinical trial in 652 study participants (Mannick et al., 2021). While a larger Phase 3 trial did not reach significance for reduction in mild RTIs, there was a clear trend to improved immune function (Mannick et al., 2021). mTOR inhibitors therefore offer a therapeutic route to enhance ageing immune responses against viral pathogens for which we currently lack effective pharmacological interventions. However, there is a gap in our understanding of how they impact on cellular processes such as immune cell ageing, which underpins immunosenescence and subsequent organismal ageing, in an immune-unchallenged steady state.

There is accumulating evidence that DNA damage is a central driver of immune cell ageing, immunosenescence, and whole-organism ageing (Kell et al., 2023, Yousefzadeh et al., 2021a, Yousefzadeh et al., 2021b). In this study, we aimed to determine whether low-dose mTOR inhibition could enhance DNA stability in human T cells, a key immune cell type affected by age-related DNA damage. Using a combination of *in vitro* DNA damage assays, *ex vivo* profiling of age-related immune cells, and a placebo-controlled *in vivo* intervention with rapamycin in older people, we sought to explore mTOR inhibitors as a potential strategy to protect cells from DNA damage and limit senescence. Our findings have implications for geriatric medicine, radioprotection during cancer therapy, and safeguarding astronauts from cosmic radiation.

## RESULTS

### DNA damage in T cells is associated with elevated mTORC signalling

In order to develop an *in vitro* model for DNA damage and a reliable read-out in primary human immune cells, we cultured isolated human peripheral blood mononuclear cells (PBMCs) from healthy donors with T cell activating antibodies against CD3 and CD28 for 3 days, followed by treatment with zeocin, a double-strand break (DSB) inducer, for 2 hours (DSBs in circulating leukocytes are predictive of increased mortality in in humans (Bonassi et al., 2021)). Acute 15-minute exposure to hydrogen peroxide was used as a positive control for DNA damage induction (**Figure 1a**). After recovery, cells were analysed by flow cytometry to identify CD4^+^ and CD8^+^ T cells, and assessed for levels of the DNA damage marker, γH2AX (gating strategy in **Figure S1**).

**Figure 1.**
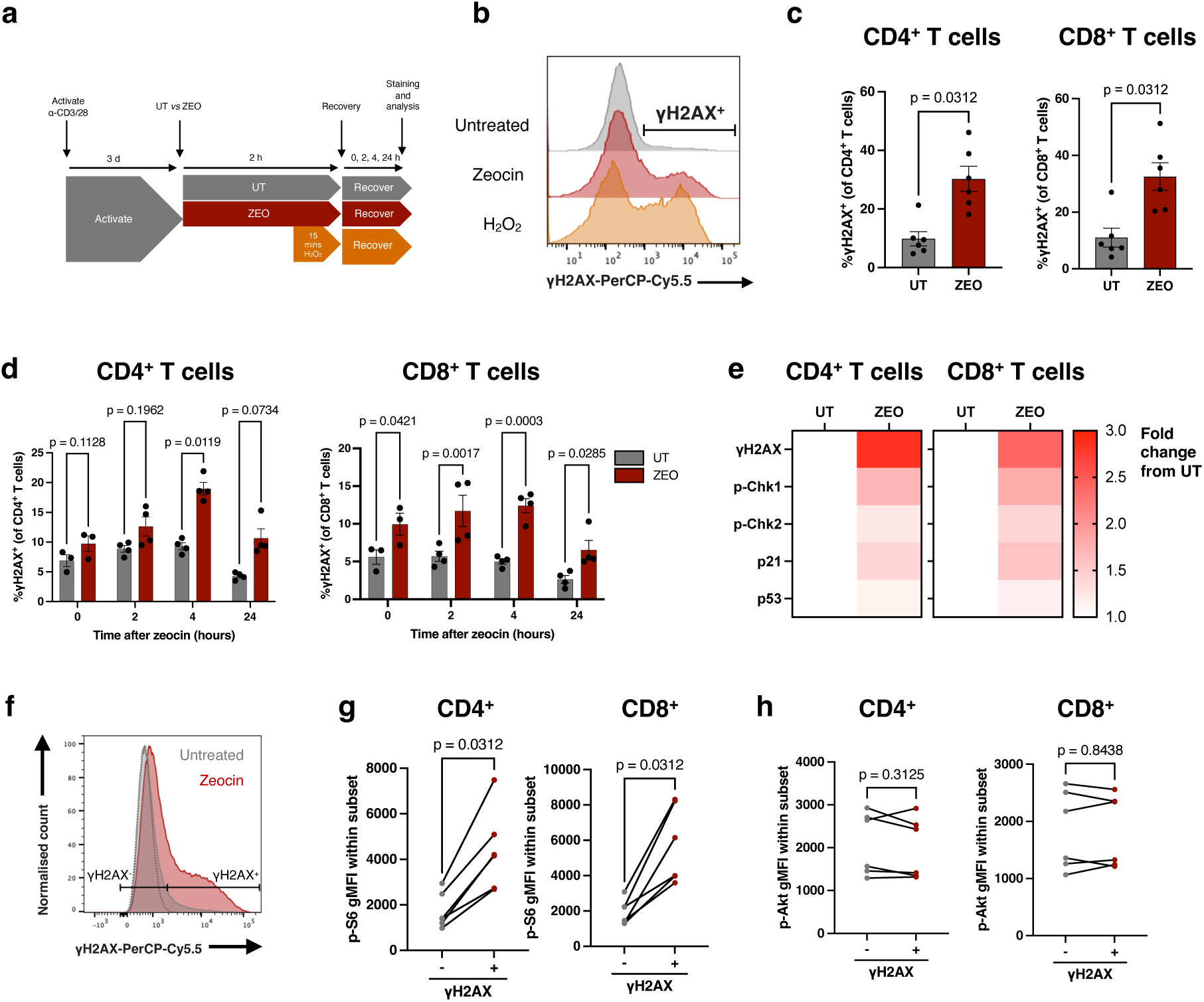
T cell DNA damage is associated with elevated mTORC1 activity. (a) DNA damage assay design. (b) Representative histograms of γH2AX levels by flow cytometry in untreated, zeocin-treated (200 µg/ml), and H_2_O_2_-treated (25 µM) PBMCs gated on CD4^+^ T cells. (c-d) Proportion of γH2AX^+^ of CD4^+^ (left) and CD8^+^ (right) T cells after (c) 4 hours recovery from zeocin treatment, n=6 healthy donors, or (d) across different recovery times after zeocin treatment, from 4 independent experiments using PBMCs from 1 donor. (e) Heatmaps for levels of DDR signalling molecules expressed as fold change from untreated (UT) cells. (f) Representative gating of γH2AX^+^ and γH2AX^-^ cells based on untreated (grey), zeocin-treated (red) cells, and fluorescence minus one (FMO, dotted grey) control. (g-h) gMFI of (g) p-S6 and (h) p-Akt in zeocin-treated CD4^+^ or CD8^+^ T cells gated as either positive or negative for γH2AX, n=6 healthy donors. P-values are derived from a two-way ANOVA with Šídák’s multiple comparisons test (d), and a Wilcoxon matched-pairs signed rank test (c, g, h).

As seen in **Figure 1b**, zeocin treatment led to a marked increase in T cells positive for γH2AX, with a similar though more extensive shift to γH2AX-positivity in the peroxide-treated controls. This increased γH2AX signal was associated with a large increase in the percentage of T cells staining positive for γH2AX, from ∼10% untreated control cells to ∼30% zeocin-treated both CD4^+^ and CD8^+^ cells (**Figure 1c**). We note that the small percentage of the untreated control cells showing γH2AX positivity potentially indicates DSB formation during activation. Levels of γH2AX positivity peaked at 4 hours post zeocin treatment, reducing by 24 hours of recovery (**Figure 1d**). Consistent with elevated γH2AX, zeocin-treated cells also showed elevated DNA damage response signalling, including phosphorylation of checkpoint kinases Chk1 and Chk2, as well as increased protein levels of tumour suppressor p53, which is stabilised by phosphorylation during the DDR, and its transcriptional target, the cyclin-kinase inhibitor p21 (**Figure 1e**). To determine whether DNA damage correlates with changes in mTOR activity, we further analysed T cells with low and high levels of γH2AX for their level of phosphorylated mTORC1 target S6 (p-S6) and mTORC2 target Akt (p-Akt) (**Figure 1f**). Notably, both CD4^+^ and CD8^+^ cells with high γH2AX signals showed significant increases in phosphorylated S6 (**Figure 1g**), an indirect target of mTORC1, but no change in levels of in mTORC2 target p-Akt (**Figure 1h**), suggesting that DNA damage in T cells is associated with elevated mTORC1 activity.

### Suppression of mTORC signalling reduces markers of DNA damage in human T cells in vitro

To test the association between high levels of DNA damage markers and elevated mTORC signalling, we assessed the impact of mTORC inhibitors on the DDR in the zeocin-induced DNA damage model. T cells were incubated throughout their 3-day activation, 2-hour zeocin-treatment, and 4-hour recovery periods with low dose mTORC1 inhibitor rapamycin (10nM), pan-mTOR inhibitor AZD8055 (100nM) or DMSO vehicle control (**Figure 2a**). Exposure to rapamycin and AZD8055 over this 3-day activation significantly suppressed p-S6 levels, apparent as early as 6 hours (**Figure S2a-b**). CD25 upregulation, a marker of T cell activation, was not impacted by mTOR inhibition at this low dose (**Figure S2c**). As before (**Figure 1b-c**), zeocin treatment resulted in a significant increase in overall γH2AX levels. However, this surge in γH2AX was greatly attenuated by treatment with the mTOR inhibitors rapamycin or AZD8055 (**Figure 2b**), reflected by the percentage of CD4^+^ T cells staining positive for γH2AX after zeocin treatment being significantly reduced by both mTOR inhibitors (**Figure 2b**, middle). By contrast, in CD8^+^ cells, this reduction was only significant on rapamycin treatment (**Figure 2b**, right).

**Figure 2.**
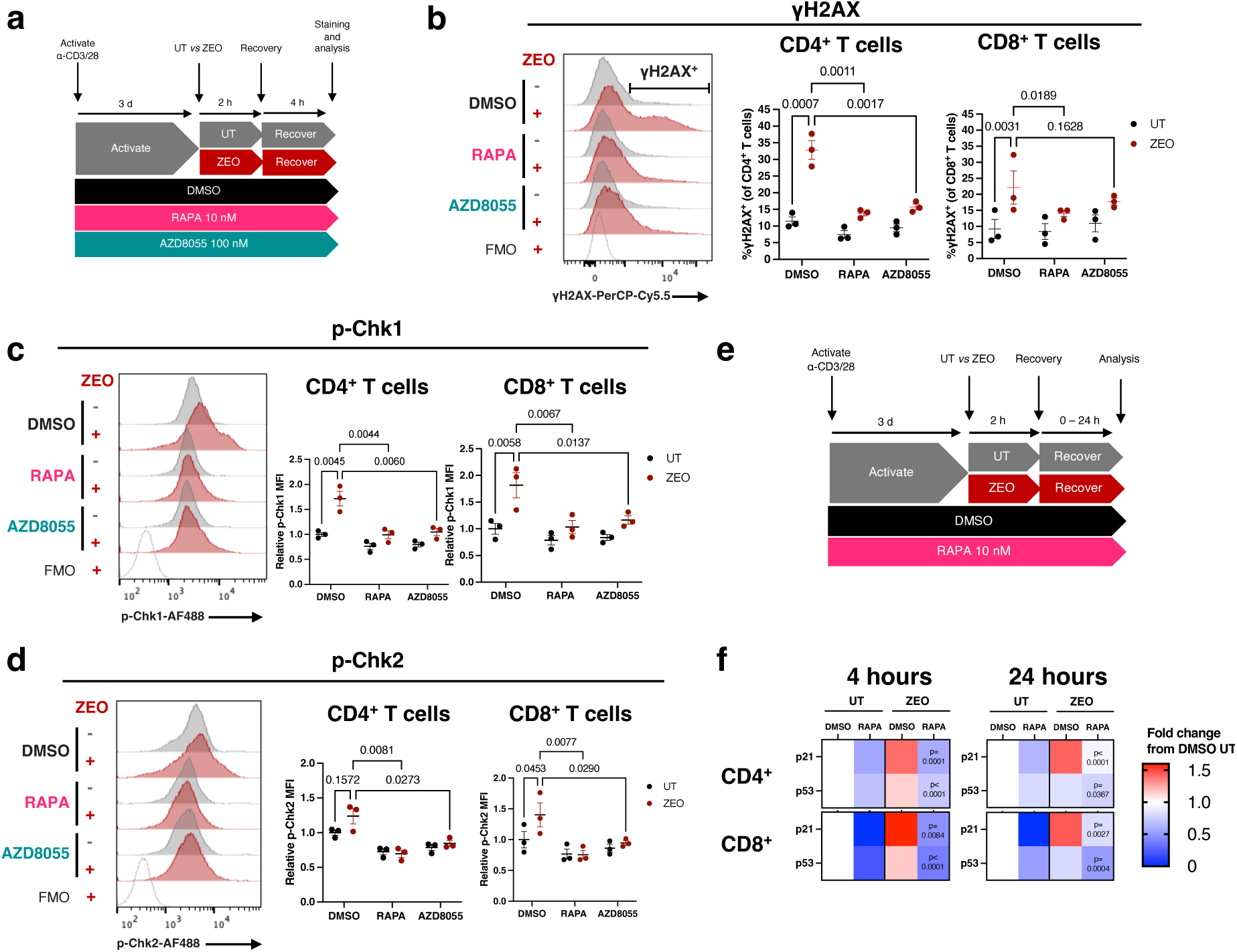
mTOR inhibition reduces the DNA damage response in genotoxin-exposed human T cells from healthy donors. (a) Experimental design. (b) Representative flow cytometry fluorescence histograms of γH2AX levels in CD4^+^ T cells with fluorescence minus one (FMO) control (left), with quantification of the proportion of γH2AX^+^ cells across conditions in CD4^+^ and CD8^+^ T cells, n=3 healthy donors. (c-d) Representative fluorescence histograms of p-Chk1 (c) or p-Chk2 (d) levels in CD4^+^ T cells, with quantification of fluorescence relative to average of DMSO untreated control in CD4^+^ and CD8^+^ T cells, n=3 healthy donors. (e) Experimental design for (f). (f) Heatmaps of gMFI of p21 and p53 assessed by flow cytometry, in CD4^+^ and CD8^+^ T cells at 4 and 24 hours recovery from zeocin, expressed as fold change from the DMSO untreated (UT) condition for each recovery time point. P-values represent comparisons to the DMSO zeocin (ZEO) condition, n=4 healthy donors. P-values are determined from a two-way ANOVA with Šídák’s (b-d) or Dunnett’s (f) multiple comparisons test.

To investigate further whether mTOR inhibition affected signalling within the DNA damage response, we assessed levels of phosphorylated (i.e. activated) checkpoint kinases Chk1 and Chk2. Both rapamycin and AZD8055 treatment prevented the zeocin-induced increase in levels for both p-Chk1 and p-Chk2 in both CD4^+^ and CD8^+^ cells (**Figure 2c-d**). We additionally assessed levels of the DDR proteins p53 and p21 at both 4 hours recovery from zeocin, and at a later 24-hour timepoint, to assess longer-term effects on resolution of the DDR (**Figure 2e**). Notably, in control cells without zeocin-induced DNA damage, rapamycin treatment led to reduction in p53 and p21 levels, compared with DMSO vehicle controls (**Figure 2f**). p21 levels increased by 4 hours recovery following zeocin treatment, remaining elevated at 24 hours; this response was completely ablated on mTOR inhibition by rapamycin treatment (**Figure 2f**). Similarly, the elevated p53 signal seen at 4 hours was significantly reduced on rapamycin treatment. By 24 hours, the p53 signal was reduced in zeocin-treated cells (with and without rapamycin treatment) compared with levels at 4 hours post damage in both CD4^+^ and CD8^+^ T cells, though mTORC inhibition led to a further significant drop in p53 levels (**Figure 2f**).

### Direct association between high levels of damage and elevated mTORC signalling

Having identified that continuous mTOR inhibition suppressed DDR upregulation, we next investigated the temporal nature of this effect, by incubating cells with rapamycin either before, during or after DNA damage by zeocin exposure (**Figure 3a**). To do this, T cells within PBMC cultures from healthy donors were activated for 3 days with anti-CD3 and anti-CD28 antibodies, then sequentially split into aliquots and incubated with rapamycin or DMSO ± zeocin as shown in **Figure 3a**. Following the recovery period also in the presence or absence of rapamycin, the percentage of cells staining positive for DNA damage marker γH2AX was assessed by flow cytometry. In all cases, zeocin treatment resulted in an increase in γH2AX-positive cells (**Figure 3b-c**), but CD4^+^ T cells showed a significant reduction of γH2AX positivity when treated with rapamycin before, during or after zeocin treatment compared with the DMSO-only controls with zeocin (**Figure 3b**). Furthermore, the fold increase in γH2AX^+^ cells significantly correlated with levels of both p-S6 and p-Akt, consistent with a role for mTOR activity in a highly DNA-damaged phenotype (**Figure 3d-e**). The exception to this was p-Akt in CD8^+^ T cells, which negatively correlated with levels of γH2AX (**Figure 3e**, right), consistent with the result that treatment with rapamycin only during or after zeocin treatment (at times when p-Akt was not suppressed) could minimise γH2AX induction. In summary, these data suggest that rapamycin treatment either before, during, or after exposure to a genotoxin, i.e. even short-term treatment could prevent zeocin-induced γH2AX levels.

**Figure 3.**
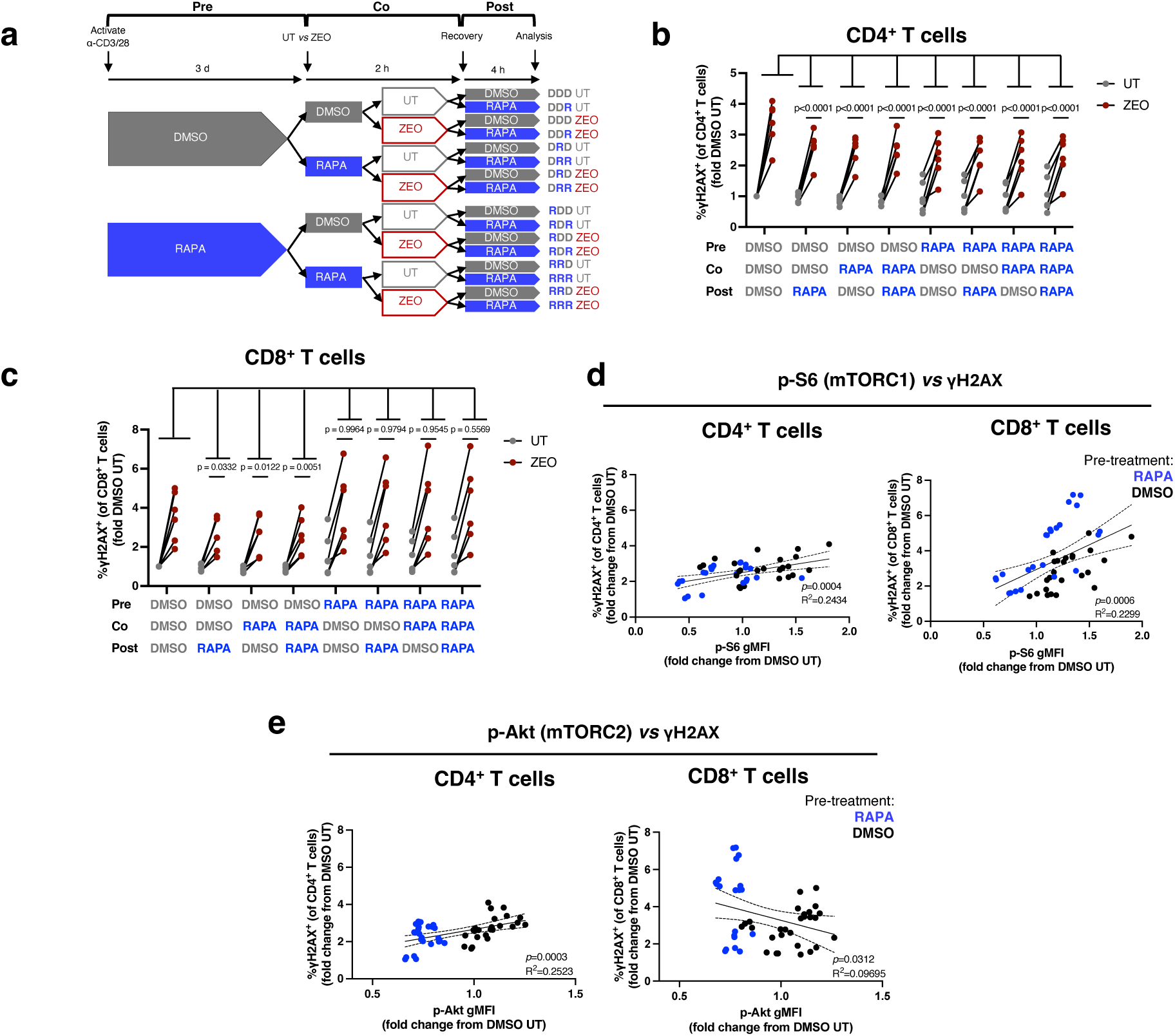
Rapamycin inhibits γH2AX at any point with respect to genotoxic treatment and suppression of mTOR activity correlates with DNA damage. (a) Experimental design. (b-c) Proportion of γH2AX+ of (b) CD4+ and (c) CD8+ T cells, in PBMCs treated with 10 nM RAPA or DMSO vehicle control pre-, co-, and/or post-treatment with zeocin. Data are expressed as fold change from DMSO→DMSO→DMSO (DDD) untreated controls for each donor. P-values represent comparisons to the DDD ZEO condition, n=6 healthy donors. (d-e) Simple linear regression analyses from experiment in (a), of γH2AX and either p-S6 (d) or p-Akt (e) in zeocin-treated CD4+ or CD8+ T cells. Values are expressed as fold change from DMSO→DMSO→DMSO UT with line of best fit and 95% confidence intervals indicated. Data are pooled from 6 independent experiments, n=6 healthy donors. P-values are derived from a two-way ANOVA with Šídák’s (b-c) multiple comparisons test.

### Reduction in DNA damage markers by mTOR inhibition is not due to impacts on cell cycle or protein synthesis

Progression through the cell cycle is halted during the initial stages of the DDR to allow for repair of DNA lesions. Thus, one possible explanation for our observation that mTOR inhibition limits γH2AX^+^ DNA damage in T cells is that it promotes cell cycle arrest to support DNA repair. We therefore first measured the proportion of cells in each phase of the cell cycle (G_0_/G_1_, S, or G_2_/M) in untreated and zeocin-treated T cells using flow cytometry (**Figure 4a**). Zeocin treatment of both CD4^+^ and CD8^+^ T cells halved the proportion of cells in S-phase (from 52% without zeocin to 26% with zeocin treatment) with a concomitant increase in the proportion of cells in G_2_/M phase (from 1.3% to 15%) (**Figure 4b**), suggesting that DNA-damaged cells proceed to G_2_ but then activate cell cycle checkpoints to prevent cell division.

**Figure 4.**
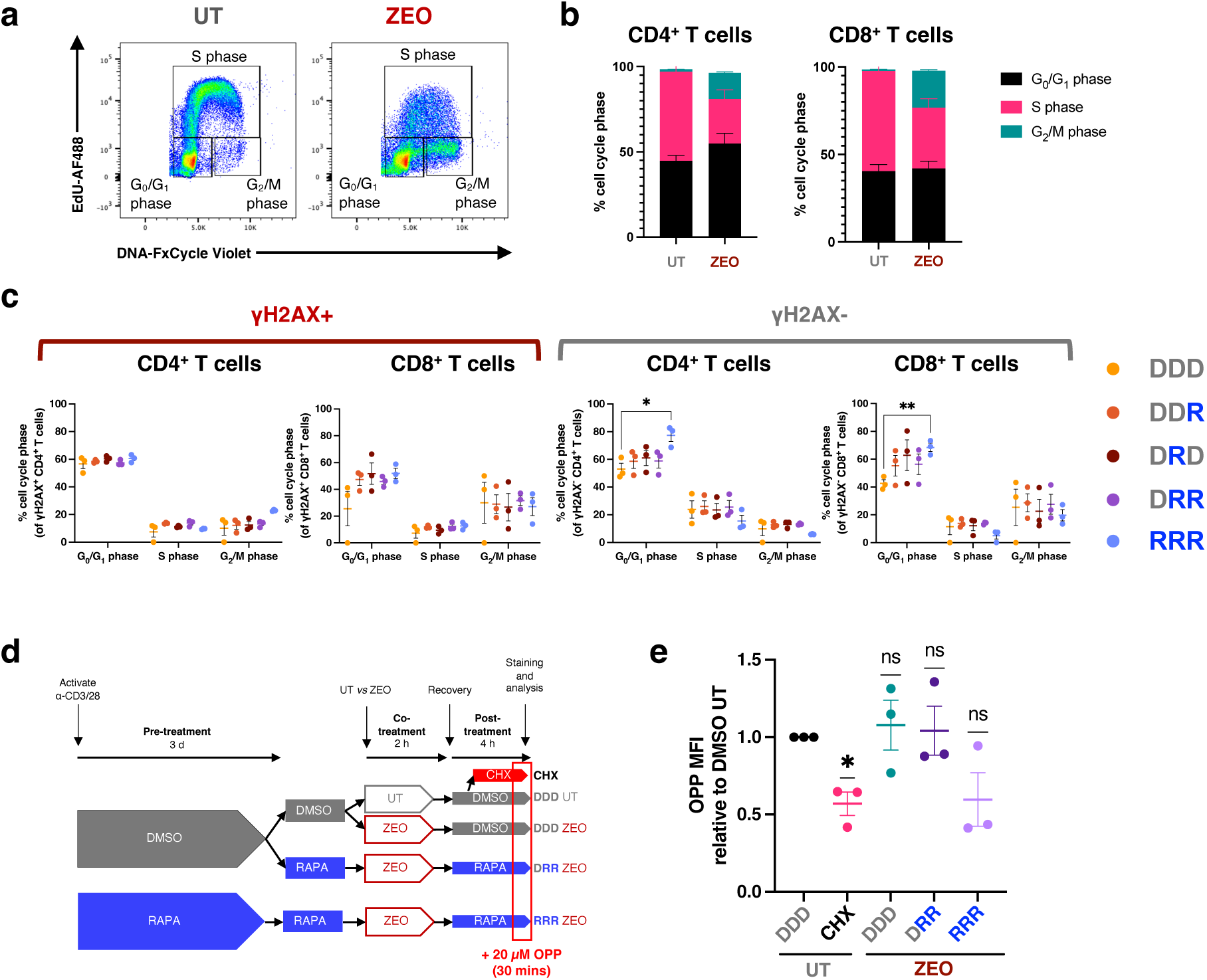
Mitigation of DNA damage markers by rapamycin is not due to modulation of the cell cycle or protein synthesis. (a-b) Representative histograms for cell cycle phases in untreated (UT) and zeocin-treated conditions with quantification IN (b), n=3 healthy donors. (c) Cell cycle phases in zeocin-treated T cells treated with rapamycin continuously (“RRR”), during (“DRD”), and/or after (“DRR”, “DDR”) zeocin exposure, gated as positive or negative for γH2AX, n=3 healthy donors. Unless otherwise indicated, statistical comparisons between different treatment conditions within each cell cycle phase (G0/G1, S, G2/M) were not significant, as determined by a two-way ANOVA with Dunnett’s multiple comparisons test. (d) Experimental design for measuring nascent protein synthesis with 20 µM O-propargyl-puromycin incorporation over 30 minutes. 1-hour treatment with 50 µg/ml cycloheximide (CHX) was used a positive control for inhibition of protein synthesis. (e) OPP mean fluorescence intensity (MFI) of CD4^+^ T cells, relative the DMSO untreated (UT) control, across conditions undergoing treatment with 10 nM rapamycin continuously (”RRR”) or during and after zeocin exposure (“DRR”). *P*-values are derived from a one-sample t-test (theoretical mean = 1), n=3 donors.

To test whether mTOR inhibition affected cell cycle progression, cells were treated with rapamycin continuously (RRR), during (DRD), and/or after treatment (DRR, DDR) with zeocin. Under these conditions previously, rapamycin limited zeocin-induced γH2AX levels in CD4^+^ and CD8^+^ T cells (**Figure 3b-c**). We observed an increase in the G_0_/G_1_-phase population on continuous rapamycin treatment (RRR) in cells without overt DNA damage (i.e. γH2AX negative), though it did not affect the proportion of G_0_/G_1_-phase cells in the γH2AX-positive population, indicating that continuous rapamycin treatment did not change cell cycle phase distribution in the context of DNA damage (**Figure 4c**). Since rapamycin treatment before, during, after zeocin treatment, or continuous exposure (DDR, DRD, DRR and RRR) effectively limited the induction of γH2AX in T cells (**Figure 3b-c**), but did not affect cell cycle phase distribution in DNA-damaged γH2AX^+^ cells (**Figure 4c**), we concluded that the effect of rapamycin on γH2AX was not due to effects on the cell cycle.

mTOR is a master anabolic regulator of protein synthesis (e.g. by activating ribosomal S6 protein through S6K-dependent phosphorylation), so it is conceivable that the reduced levels of DNA damage proteins we detect by flow cytometry may be a consequence of blockade of their *de novo* synthesis (albeit that the acute DDR is predominantly mediated post-translationally). To evaluate the effects of rapamycin on nascent protein synthesis in the DNA damage assay, cells were treated with rapamycin or DMSO vehicle control before, during and/or after zeocin treatment and then incubated for the final 30 minutes of their 4-hour recovery from zeocin with O-propargyl-puromycin (OPP), an alkyne analogue of puromycin that is incorporated into nascent polypeptides and halts further translation (**Figure 4d**). The mean fluorescence of labelled OPP in cells thus reports short term total *de novo* protein synthesis. One-hour treatment with 50 μg/ml cycloheximide (CHX) served as a positive control for inhibition of protein synthesis. As expected, CHX-treated cells incorporated significantly less OPP than the DMSO untreated controls (**Figure 4e**). Rapamycin treatment both during and after zeocin exposure (DRR) did not significantly affect OPP incorporation, though continuous rapamycin treatment (RRR) showed a non-significant trend towards lower OPP fluorescence (**Figure 4e**). Since there was no consistent effect of rapamycin in decreasing OPP levels, this suggests that the effect of rapamycin on limiting zeocin-induced DDR signalling levels was not due to decreasing global protein synthesis. In summary so far, our data indicates that the rapamycin-mediated protection from upregulation of the DDR following zeocin exposure is likely independent of effects on the cell cycle and protein synthesis.

Autophagy is required to limit DNA damage in T cells, but rapamycin’s protective effect is autophagy-independent.

Autophagy is a cytoprotective cell recycling process that is repressed by mTORC1 activity; notably, autophagy is also involved in regulation of the DNA damage response (Vessoni et al., 2013). We therefore probed whether the mechanism by which rapamycin treatment reduces markers of DNA damage signalling following zeocin exposure, could be due to enhancement of autophagic flux, as measured by a flow cytometry-based LC3 assay (**Figure S3a-b**) (Alsaleh et al., 2020). In the presence of zeocin-induced damage (**Figure 5a**), activated T cells with a high DNA damage load (i.e. positive for γH2AX) showed significantly lower autophagic flux than those cells negative for γH2AX (**Figure 5b**). This suggests either that cells bearing a heavy DNA lesional load are less able to undergo autophagy, or that those with effective autophagy rapidly resolve DNA damage leading to low levels of damage markers such as γH2AX.

**Figure 5.**
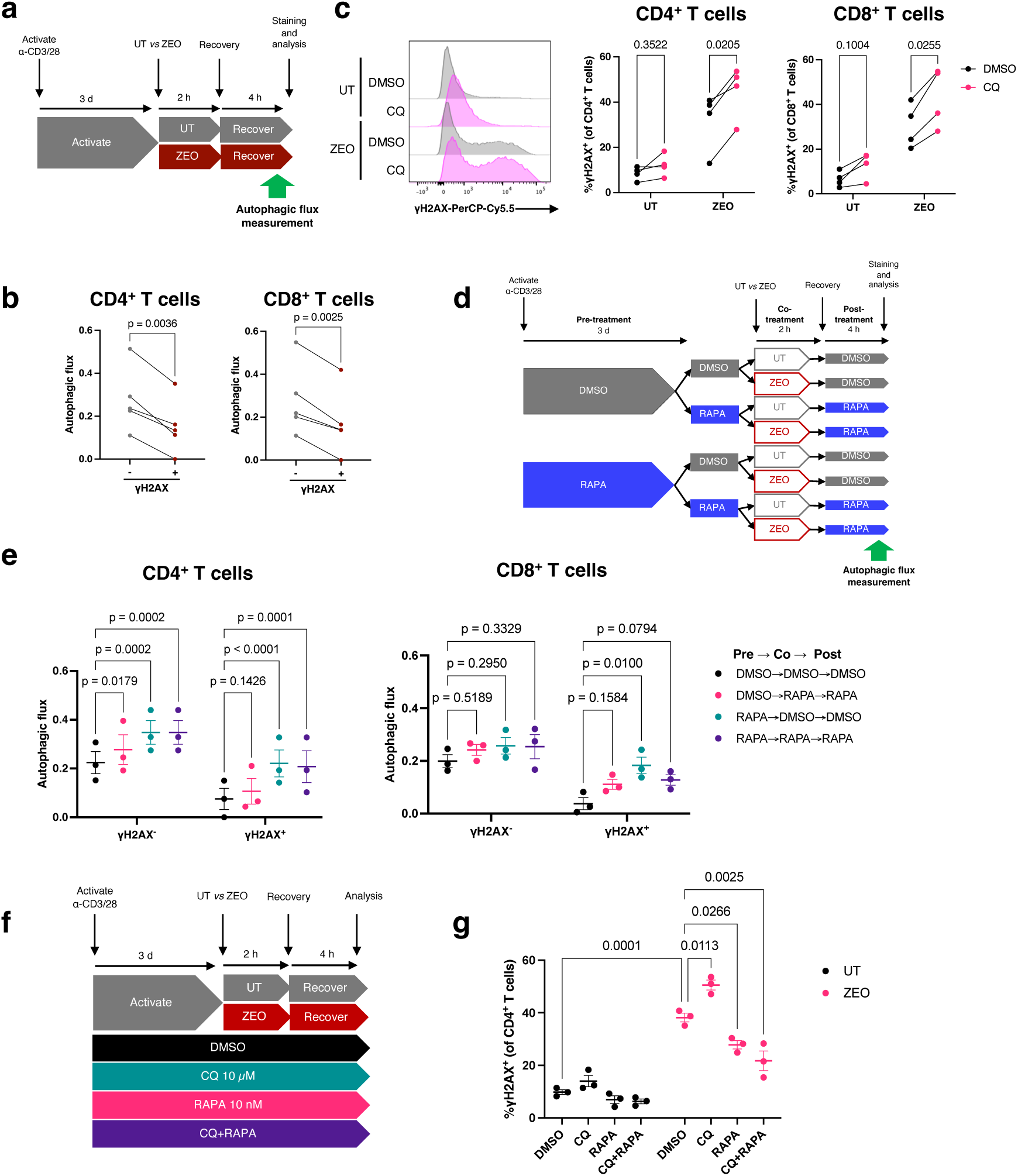
Autophagy is required for the resolution of DNA damage, but is not required for the mitigation of DDR upregulation by rapamycin. (a) Experimental design for combined DNA damage and autophagic flux assay. (b) Quantification of autophagic flux in γH2AX^+^ and γH2AX^-^ populations in flow cytometry-gated CD4^+^ and CD8^+^ T cells, n=5 healthy donors. (c) Representative γH2AX histograms (left) and quantified percentage (right) of γH2AX^+^ CD4^+^ and CD8^+^ T cells after zeocin treatment following 3-day T cell-specific activation in chloroquine (CQ, 10 µM) or DMSO control, n=4 healthy donors. (d) Experimental design for measuring autophagic flux in CD4^+^ and CD8^+^ T cells across different rapamycin treatment conditions with respect to zeocin exposure. (e) Autophagic flux in zeocin-exposed cells gated as positive or negative for γH2AX, across rapamycin treatment conditions. (f) Experimental design for treatment of zeocin-exposed activated T cells with chloroquine (CQ), rapamycin (RAPA), or dual treatment (CQ+RAPA). (g) Percentage of γH2AX^+^ CD4^+^ T cells after zeocin treatment across conditions in (f), n=3 healthy donors. P-values are derived from a paired t-test (b), a two-way ANOVA with Šídák’s (d,e) or Tukey’s multiple comparisons test (g).

To distinguish between these possibilities, we used the drug chloroquine to inhibit autophagy, which effectively halved autophagic flux in activated T cells (**Figure S3c**). In zeocin-treated cells, autophagy blockade increased γH2AX-positive cells, confirming that autophagy does limit DNA damage in human T cells (**Figure 5c**). Next, we asked whether rapamycin enhanced autophagy in zeocin-exposed cells (**Figure 5d**) and found that this was the case, regardless of γH2AX levels or timing of rapamycin administration (**Figure 5e**). We then asked whether rapamycin’s protective effect on DNA damage depended on autophagy by co-treating cells with rapamycin and chloroquine (**Figure 5f**). As expected, chloroquine inhibited autophagic flux and increased γH2AX positivity, but rapamycin still markedly reduced DNA damage despite strong autophagy inhibition in the context of chloroquine co-treatment (**Figure S3d, Figure 5h**). These findings indicate that while autophagy supports DNA damage resolution, rapamycin’s protective effect is independent of cell cycle pausing, protein synthesis, and autophagy.

### Rapamycin decreases overall DNA lesional burden and reduces T cell death following DNA damage

DDR signalling requires activation of several PI3-like kinases (e.g. DNA-PKcs, ATM and ATR). It is therefore possible that mTOR inhibitors reduce apparent DNA damage by inhibiting critical DDR enzyme signalling, in a manner that would be highly detrimental to cell health and survival. Alternatively, reduced levels of DDR signalling may instead reflect a lower DNA lesional burden. To distinguish between these two possibilities, we assessed the extent of DNA breaks after 4 hours of recovery from zeocin exposure, using the alkaline comet assay, in isolated CD4^+^ T cells treated with or without rapamycin (**Figure 6a**). Treatment with hydrogen peroxide was used as a positive control for DNA breakage. Both zeocin and hydrogen peroxide treatment significantly increased DNA lesions (both DSBs and SSBs) compared to untreated controls (**Figure 6b-c**). Notably, DNA lesion burden was markedly reduced in CD4^+^ T cells treated with rapamycin at this 4-hour recovery timepoint from zeocin. This suggests that the reduction in DDR signalling afforded by rapamycin is due to enhance genome stability rather than downstream inhibition of DDR enzymes (**Figure 6b-c**).

**Figure 6.**
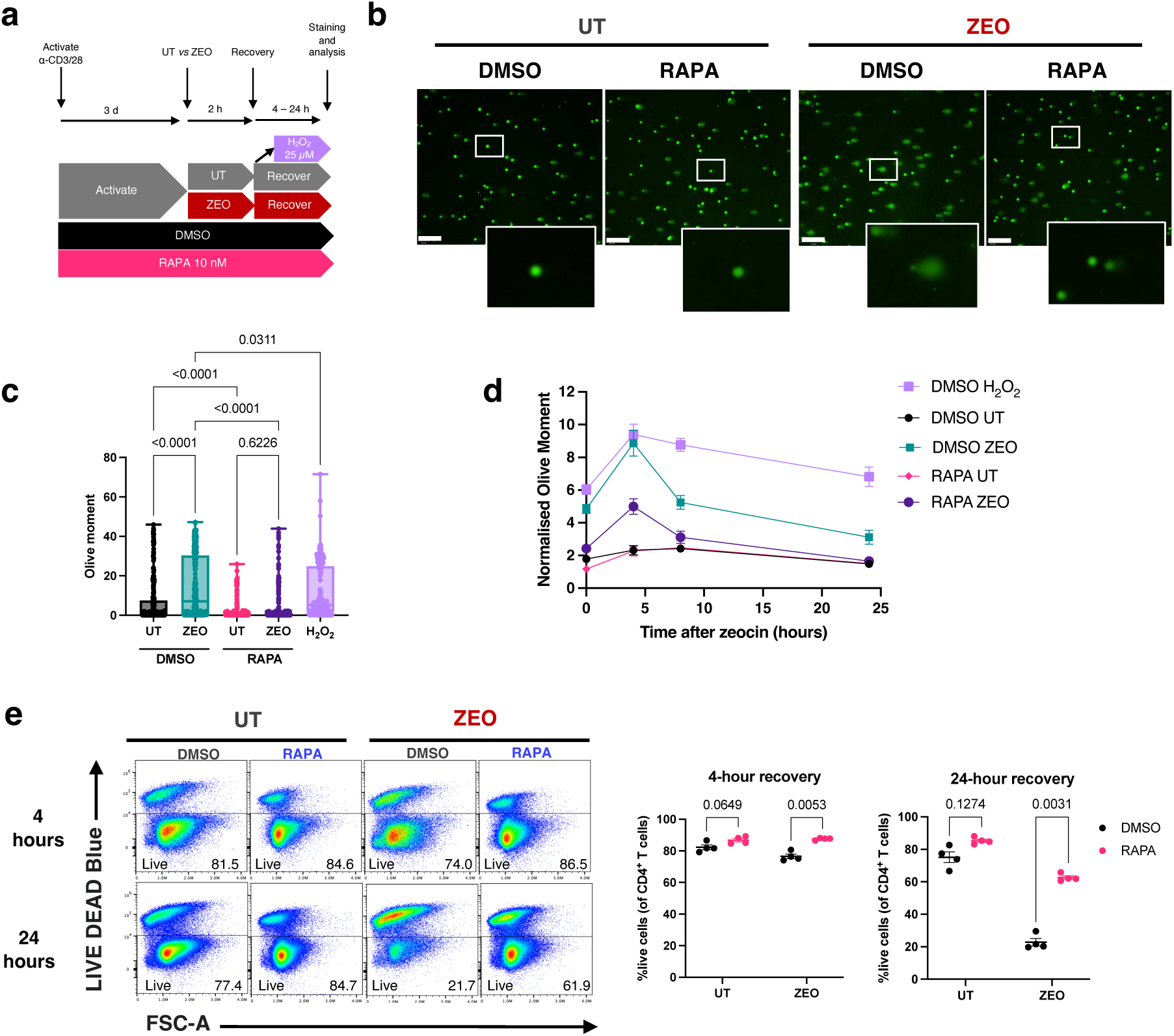
Rapamycin attenuates DNA lesional burden and improves survival after exposure to a DNA-damaging agent. (a) Experimental design for zeocin treatment (200 µg/ml) following continuous exposure to rapamycin (10 nM) or DMSO vehicle control in isolated CD4^+^ T cells. (b) Representative images of comets, scalebar represents 200 µm. (c) DNA lesions as measured by the Olive moment of >250 comets analysed across conditions following a 4-hour recovery from zeocin. (d) Comet Olive moments throughout a 24-hour time course of recovery from zeocin normalised to the median value of the DMSO untreated (UT) condition at each time point. Data are the median ± SEM of 100-250 nuclei analysed per condition. Representative of three independent experiments, n=3 healthy donors. (e) Live cells across conditions, as measured by lack of fluorescence of a membrane-permeable dye, at 4 and 24 hours recovery from zeocin exposure. P-values are derived from a one-way ANOVA with Tukey’s multiple comparisons test (c), or a two-way ANOVA with Šídák’s multiple comparisons test (e).

To assess further the kinetics of DNA lesional burden and potential resolution of damage, we then assessed comet Olive moment in cells incubated continuously with rapamycin over a time course of up to 24 hours after exposure to zeocin. The peak of DNA lesions manifesting as comet tails was found to occur at 4 hours after zeocin treatment (**Figure 6d**). Continuous rapamycin treatment significantly limited the DNA lesion burden at all timepoints tested (**Figure 6d**). Notably, rapamycin reduced comet tails even at 0 hours post-zeocin exposure, i.e. directly after genotoxin treatment, suggesting a stark enhancement of resilience from DNA damage that may reflect prevention of DNA lesion formation. In summary, we can rule out a negative effect of rapamycin inhibiting PI3-like kinases in the DDR, and instead propose that rapamycin positively protects cells from DNA damage.

To explore whether this effect of rapamycin on reducing lesional load has an impact on overall cell physiology, we measured cell viability by assessing fluorescence of a membrane-impermeable dye that is taken up only by dead cells (**Figure 6e**). Consistent with the increase in cells with major DNA lesions following zeocin exposure, we observed a decrease in the percentage of live CD4^+^ T cells at 4 hours, leading to a severe reduction to only 20% live cells by 24 hours recovery from zeocin in DMSO vehicle control cells i.e. the high lesional burden induced by zeocin treatment is lethal to the majority of T cells (**Figure 6e**). Remarkably, continuous treatment with low-dose rapamycin (10 nM) supported much greater cell survival with over 60% cells still viable 24 hours after zeocin treatment (**Figure 6e**), strongly suggesting that rapamycin does indeed enhance DNA repair responses i.e. it may act as a genoprotector.

### Age-related immune subsets show elevated markers of DNA damage, cell senescence, and mTOR activity

These findings suggest a genoprotective role for rapamycin in the immune system, but are based on an *in vitro* model of acute DNA damage. In humans, immunosenescence involves the expansion of terminally-differentiated immune subsets linked to dysfunction (**Table 1**), though their exact phenotype remains unclear. Since chronic DNA damage accumulation is a key hallmark of ageing, that in immune cells contributes to whole-body ageing (Yousefzadeh et al., 2021b), we examined whether aged human immune cells show signs of DNA damage and cell senescence, and whether this correlates with their mTOR activity.

**Table 1.**
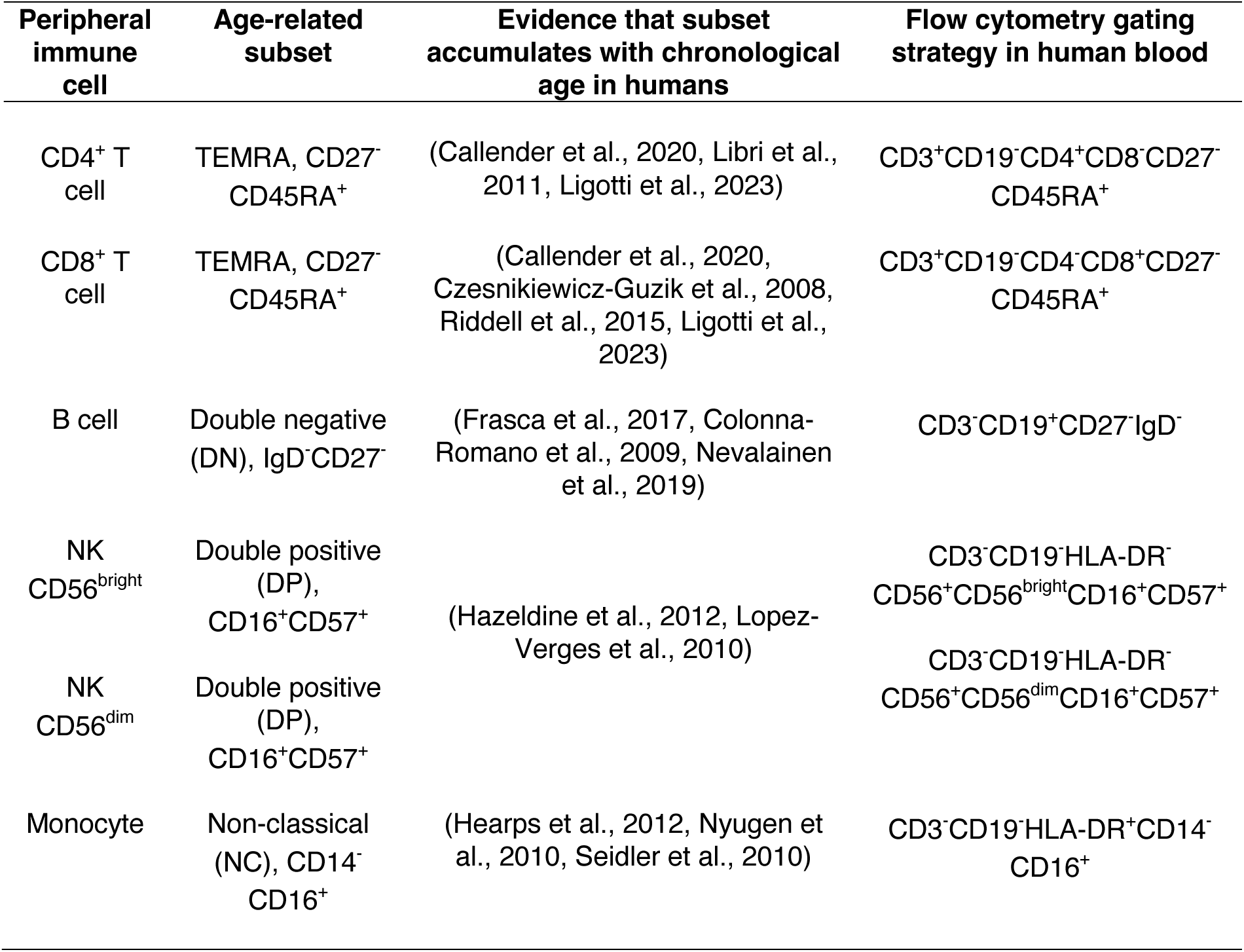
Identification of age-related peripheral immune subsets in humans.

Using 27-colour spectral flow cytometry, we analysed age-related immune cell subsets from healthy donor blood, including TEMRA T cells (CD4^+^ and CD8^+^), IgD^-^CD27^-^ (double-negative) B cells, CD16^+^CD57^+^ NK cells, and non-classical monocytes (**Figure 7a, Figure S4, Table 1**). We then assessed senescence- and DNA damage-associated markers (p21, p16, p53, γH2AX) and cell size (measured by forward scatter, FSC) to determine whether these subsets were enriched for ageing biomarkers compared with their naïve counterparts (Stein et al., 1999, Passos et al., 2007, van Deursen, 2014, Tsai et al., 2021).

**Figure 7.**
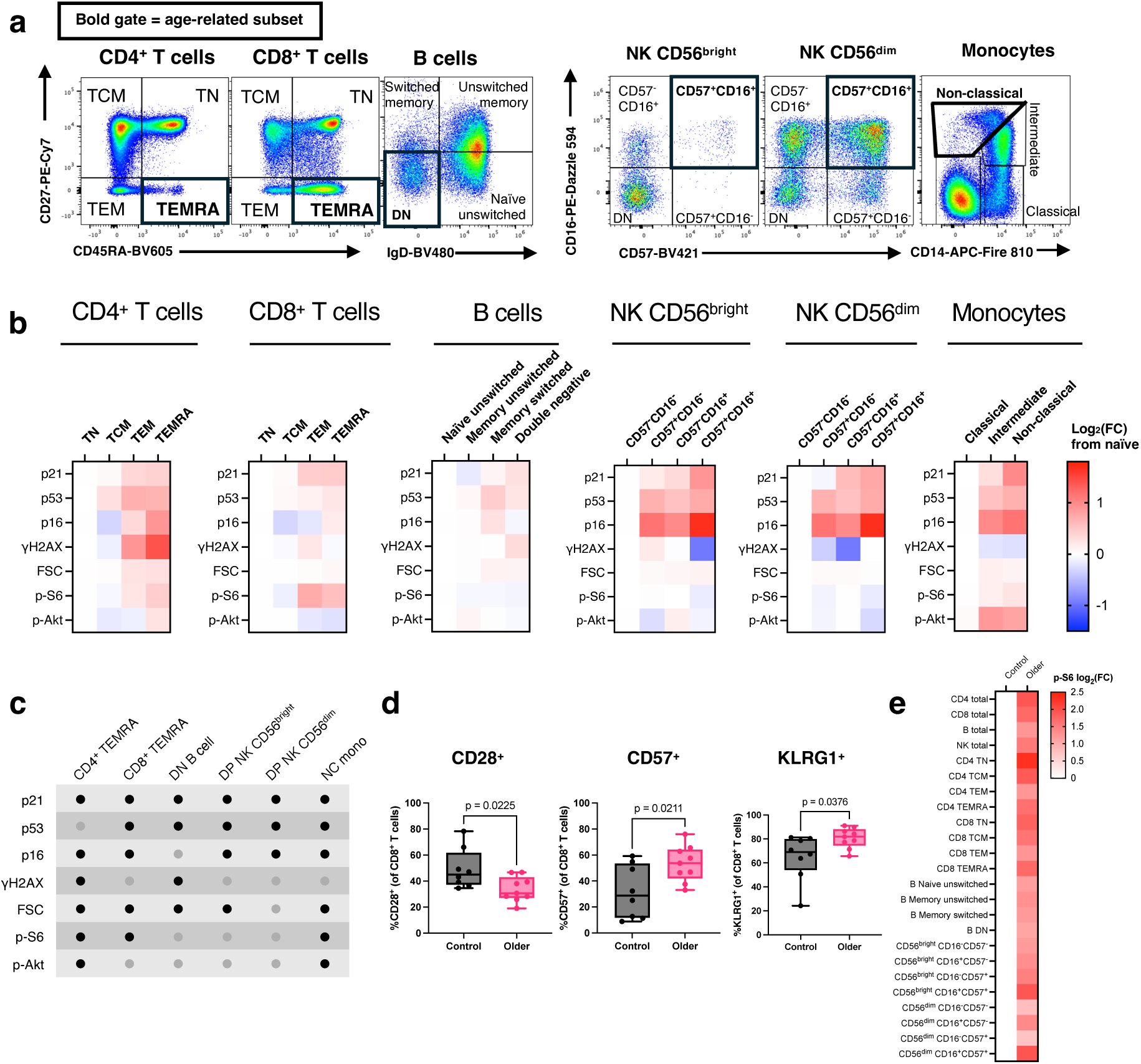
Age-related peripheral immune subsets from healthy donors display elevated markers of cellular senescence and mTORC1/2 hyperactivity. (a) Immune cell subsets in PBMCs from a healthy donor identified by flow cytometry. Gates highlighted in bold indicate age-related immune subsets. (b) Geometric mean fluorescence intensity (gMFI) of biomarkers for senescence and mTORC1/2 activity as measured by spectral flow cytometry across immune subsets in n=8 healthy donors. Data represent log2(fold change) from the mean of the lefthand, most early-differentiated immune cell population for each cell type. FSC = forward scatter. (c) Table summarising significantly increased markers (black dots) in age-related immune subsets compared to early-differentiated immune cell counterparts, as assessed with one-way ANOVAs with Dunnett’s multiple comparisons test between age-related and early-differentiated subsets for each cell type (n=8 healthy donors). (d) Proportion of CD8^+^ T cells positive for CD28, CD57, and KLRG1 in PBMCs from healthy younger donors (17-50 years old, n=8) and older donors (average age 62 years old, n=9). P-values are derived from unpaired t-tests. (e) Geometric mean fluorescence intensity (gMFI) of p-S6 in immune subsets in older donors (n=9) expressed as log2(fold change) from the mean value of cells of control volunteers (n=8), assessed by flow cytometry.

We observed that senescence markers were significantly enriched in age-related immune cells compared to their naïve equivalents, with each of the 6 subsets assessed showing significant elevation of at least 3/5 senescence markers (**Figure 7b-c**). In particular, age-related CD4^+^ and CD8^+^ TEMRAs and non-classical monocytes showed the greatest number of elevated senescence markers compared to early-differentiated cells of the same lineage (4/5 each, **Figure 7c**). Double-negative (DN) B cells and CD57^+^CD16^+^ (double-positive, DP) NK cells all exhibited significant upregulation of 3/5 senescence markers (**Figure 7c**). Notably, DNA damage marker γH2AX was elevated only in age-associated T and B cells i.e. immune cell types that undergo double strand breaks during V(D)J recombination to form T cell and B cell receptors (**Figure 7b-c**). Intriguingly, the most common senescence feature that was increased across age-related subsets was p21, present in all 6 age-related subsets compared to their early-differentiated controls (**Figure 7c**). This was followed by high p53, p16, and cell size/FSC (present in 5/6 subsets each, **Figure 7c**). Overall, these data suggest that age-related subsets display several features of cellular senescence, and particularly overexpress the p21 and p53 pathway, suggesting a DNA-damage induced senescence phenotype. Importantly, these data demonstrate that age-related immune cells may be targetable with genoprotective senotherapeutics.

Next, we asked whether these age-related subsets showed changes in their activation of mTORC1 and mTORC2 by measuring their levels of p-S6 and p-Akt respectively. We observed that age-related CD4^+^ TEMRAs, CD8^+^ TEMRAs, and non-classical monocytes all showed elevated p-S6, with CD4^+^ TEMRAs and non-classical monocytes additionally displaying increased p-Akt levels (**Figure 7b-c**). These findings indicate that while senescence markers were present in all age-related immune subsets, mTOR hyperactivation occurs only in T cell and monocyte ageing.

Given that diverse age-related subsets within healthy donors showed elevated senescence and mTORC1/2 markers, we next asked whether immune cells from older people (average age 62 years-old, n=9) exhibited increased mTORC activation compared to those from younger donors (<50 years old, n=8). We first verified that, compared to the younger group, older donors had an increased percentage of CD8^+^ T cells expressing CD57 and KLRG1, and loss-of-expression of CD28 (**Figure 7d**), all of which are markers of immunosenescence (Kell et al., 2023). Comparison of immune subsets between these two groups showed that, in addition to these markers of T cell senescence, immune ageing corresponded with an increase in p-S6 levels across all immune cell types analysed (**Figure 7e**). This suggests that mTORC1 activity is a broad biomarker of human immune ageing shared by cell types from diverse lineages.

### Low-dose rapamycin reduces markers of senescence and DNA damage in humans *in vivo*

Taken together, our data so far show that age-related immune subsets exhibit features of DNA damage, cell senescence, and mTOR hyperactivation, and that human ageing is accompanied by increased mTOR activity across all immune cell types (**Figure 7d-e**). We have further demonstrated that treatment with low-dose mTOR inhibitors improve survival and reduce markers of senescence and DNA damage in human T cells treated with a genotoxic agent outside of the body. Such findings are important but require *in vivo* data before they support further clinical action. We therefore assessed whether rapamycin treatment impacts on immune cell DNA damage and senescence *in vivo* in humans, analysing PBMCs from participants of a single-blind, placebo-controlled trial (NCT05414292), in which older male volunteers received either 1 mg/day rapamycin (n=4) or placebo (n=5) for 4 months (**Figure 8a**). While the primary endpoint was to assess changes in muscle mass and protein synthesis, we aimed to assess features of immunosenescence in PBMCs isolated at several timepoints throughout the trial.

**Figure 8.**
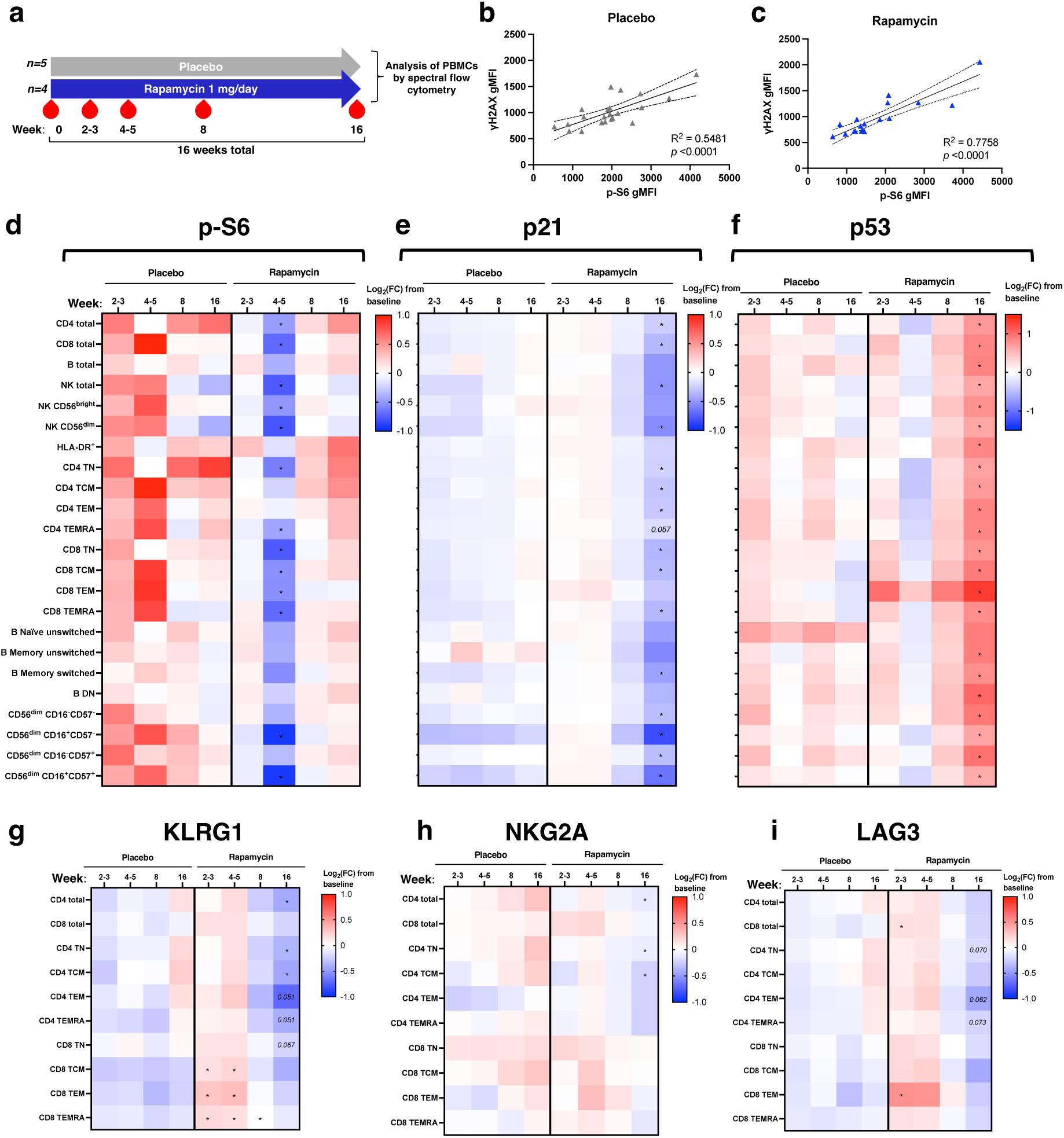
Low-dose rapamycin in vivo attenuates biomarkers of immune cell senescence and exhaustion. (a) Outline of the rapamycin experimental medicine trial. (b-c) Simple linear regression analyses of γH2AX and p-S6 geometric fluorescence intensity (gMFI) over all timepoints in CD4^+^ T cells from (b) placebo and (c) rapamycin groups. Graphs show line of best-fit with 95% confidence intervals. (d-f) Log2(fold change) from baseline in gMFI of (d) p-S6, (e) p21, and (f) p53 across immune subsets. (g-i) Percentage of defined T cell subsets positive for (g) KLRG1, (h) NKG2A, and (i) LAG3 in participants. In (d-e), each value is expressed as log2(fold change) from baseline for each participant. P-values are derived from an unpaired t-test between placebo (n=5) and rapamycin (n=4) at each time point. Statistically significant (*p<0.05) p-values are indicated.

Participants in the rapamycin and placebo groups were well-matched for age and BMI (**Table 2**). After 8 weeks of intervention, the concentration of rapamycin in the blood reached an average of 3.24 ± 1.81 nM in the treatment group (**Figure S5a**), i.e., within the same order of magnitude as the doses used in our *in vitro* experiments (10 nM). To address concerns of immunosuppression by rapamycin, the white blood cell count was assessed at 8 weeks; there were no significant differences in leukocyte counts in the blood over the initial 8-week rapamycin treatment period, and between rapamycin treated and placebo controls, suggesting that this low-dose rapamycin treatment regimen was not immunosuppressive (**Figure S5b**). To assess whether mTOR activity was inhibited at this low dose of rapamycin, we analysed p-S6 levels across immune subsets. We observed a significant decrease in p-S6 levels in most immune subsets in the rapamycin-treated participants compared to those in the placebo group at 4-5 weeks, suggesting successful inhibition of mTORC1 (**Figure 8d**). Taken together, low-dose rapamycin treatment led to detectable stable blood rapamycin concentrations at a level well below that used therapeutically for immunosuppression with no evidence of leukocyte suppression, plus reduced markers of mTORC1 activity in peripheral immune cells after 4-5 weeks.

**Table 2.** Baseline characteristics of participants in the Rapamycin trial.

|  | **Placebo (n=5)** | **Rapamycin (n=4)** | **p-value** |
| --- | --- | --- | --- |
| **Age** (years) | 64.2 ± 5.34 | 60.0 ± 4.62 | 0.1667 (ns) |
| **BMI** (kg/m<sup>2</sup>) | 26.4 ± 3.31 | 27.1 ± 1.24 | >0.9999 (ns) |

Data are presented as mean ± SD. *P*-values are derived from Mann-Whitney tests. ns = not significant.

To determine whether features of immunosenescence were impacted by *in vivo* rapamycin treatment, we analysed PBMCs from the trial using 27-colour spectral flow cytometry (**Figure S4**). Simple linear regression analyses in circulating CD4^+^ T cells revealed a strong and highly significant positive correlation between p-S6 and γH2AX levels in both treatment groups, indicating that mTOR activity and DNA damage are positively linked *in vivo* (**Figure 8b-c**). Notably, T cells from the rapamycin group showed lower p-S6 levels than those from the placebo group, which corresponded with decreased γH2AX levels (**Figure 8c**). Overall, rapamycin treatment led to a trend towards lower γH2AX levels in immune subsets, particularly in age-related CD4 TEMRA and double-negative B cells, which with higher participant numbers might show significance (**Figure S5c**). Consistent with these positive effects on γH2AX-marked DNA damage, 4-month rapamycin treatment caused a robust and significant decrease in p21 expression across most immune cell subsets studied, reflecting the attenuation of DNA damage-induced p21 with rapamycin we observed *in vitro* (**Figure 8e**). p53 expression was elevated at 4 months in PBMCs from the rapamycin-treated compared to the placebo groups (**Figure 8f**). A previous study demonstrated that *in vivo* mTOR inhibition decreased the percentage of circulating PD-1^+^ T cells (Mannick et al., 2014). Though we did not observe changes in PD-1 in the current trial (**Figure S5d**), the proportion of T cells expressing other immune co-inhibitory molecules, such as KLRG1 (**Figure 8g**), NKG2A (**Figure 8h**), and LAG3 (**Figure 8i**), was reduced in the T cells from rapamycin compared to placebo groups. Overall, these results suggest that rapamycin reduces the expression of immune exhaustion markers, and p21, a marker of cell senescence. While participant numbers in the rapamycin *in vivo* study are low, the changes in DNA damage and senescence markers is significant.

## DISCUSSION

While mTOR inhibition is a well-known and potent anti-ageing intervention in animal models, an explanation for its ability to extend health- and lifespan so reproducibly has been lacking (Weichhart, 2018). Furthermore, our understanding around why mTOR inhibitors have shown benefit in boosting immune resilience in older people is incomplete. In this study, we have demonstrated for the first time that mTOR inhibitors can protect T cells from DNA damage and senescence marker upregulation after exposure to a genotoxic agent. We show that this is through a mechanism independent of autophagy, cell cycle progression, and protein synthesis. Rather, we show that this is through a mitigation of DNA lesional burden, affording a greater survival following exposure to DNA-damaging treatment. This enhancement of protection from DNA damage, which we call genoprotection, offers a new explanation for previous studies that have demonstrated an attenuation of replicative senescence with mTOR inhibitors in 2D cell culture (Walters et al., 2016, Park et al., 2020, Iglesias-Bartolome et al., 2012) and *in vivo* in human skin (Chung et al., 2019), and its potent geroprotective ability. Our study also provides a novel explanation for previous reports which show that rapamycin improves aged antigen-specific immunity in mouse models of immunosenescence bearing immune-specific knockout of DNA repair (Yousefzadeh et al., 2021b). Furthermore, our findings expand on previous research showing a reduction in DNA damage markers with rapamycin in irradiated normal oral keratinocytes (Iglesias-Bartolome et al., 2012), DNA repair-deficient fibroblasts (Saha et al., 2014), human oocytes undergoing *in vitro* maturation (Yang et al., 2022), DNA repair-deficient mouse podocytes (Braun et al., 2025), and lymphocytes of kidney transplant patients (Chebel et al., 2016). This enhanced resilience to DNA damage with mTOR inhibition was shown to arise from several sources, including heightened expression of antioxidant enzymes, such as mitochondrial superoxide dismutase, that limit ROS and genotoxic stress (Iglesias-Bartolome et al., 2012), and increased protein expression of the DNA repair factors, MGMT and NDRG1, via a post-transcriptional mechanism (Dominick et al., 2017). Our data from human immune cells may therefore reflect a universal impact of rapamycin on promoting genome integrity in eukaryotes.

In the present study, we asked whether age-related whether age-related immune cells from diverse haematopoietic lineages exhibited DNA damage-induced senescence, by comprehensively profiling senescence markers in human immune subsets using high-dimensional spectral cytometry. Our data are the first to show that age-related immune subsets from diverse immune lineages, in CD4^+^ and CD8^+^ T cells, B cells, NK cells, and monocytes, are uniformly enriched for senescence biomarkers. In particular, the DNA damage-induced cyclin kinase inhibitor, p21, was a senescence marker upregulated in most age-related immune subsets. This suggests that the form of senescence which immune cells undergo with ageing may be p53- and p21-driven, hinting – importantly – towards a more DNA damage-induced type of senescence, consistent with other evidence that DNA damage plays a central role in the decline of immune system function (Kell et al., 2023). Immune cells from older donors exhibited higher levels of p-S6, indicating mTORC1 activity, suggesting that, like ageing of other human tissues (Markofski et al., 2015), immune ageing is associated with mTORC1 hyperactivity.

Most importantly, our findings translate to the *in vivo* condition in humans. Through a small pilot trial with limited participant numbers, ours is the first study demonstrating a significant reduction in p21-marked cellular senescence upon 4-month, low-dose rapamycin treatment *vs* placebo in immune cells in the blood of older people. We also found that rapamycin increased p53 levels in circulating immune cells. p53 serves multiple physiological roles *in vivo*; for example, in addition to its well-known role in signal transduction of acute DNA damage, it also regulates mitochondrial respiration – indeed, mice null for p53 have very poor exercise tolerance with early fatigue onset (Bartlett et al., 2014). Though at this stage highly speculative, it is possible that elevated p53 in immune cells from rapamycin-treated participants may indicate better overall metabolic health. In addition, enhancement of p53 expression has been shown recently to improve DNA repair after irradiation-induced senescence of human dermal fibroblasts (Miller et al., 2025). Therefore, while p53 was suppressed by rapamycin following acute DNA damage *in vitro*, our observation that longer-term rapamycin administration in older individuals increases p53 levels may reflect improved genome integrity.

Our findings allow us to speculate that the positive effect of 6-week treatment with the rapalogue RAD001 (everolimus) on boosting flu vaccine responses and respiratory infections may be through an attenuation of immune cell DNA damage and subsequent senescence (Mannick et al., 2018, Mannick et al., 2014). In the cited studies, everolimus caused a reduction in the proportion of circulating PD-1^+^ CD4^+^ and CD8^+^ T cells (Mannick et al., 2014). In our study, we observed a significant reduction in both KLRG1^+^ and NKG2A^+^ CD4^+^ T cells, and near-significantly LAG3^+^ CD4^+^ T cells in the rapamycin compared to placebo groups. Like PD-1, these three cell-surface proteins are all immune checkpoint inhibitors, each with roles in limiting T cell activation. Therefore, we observed similar functional effects of rapamycin as perhaps potentiating a less-exhausted T cell phenotype. Overall, based on our *in vitro* effects mTOR inhibitors and *in vivo* effects of rapamycin, we suggest that rapamycin positively enhances genome stability, and therefore targets a central hallmark of ageing (Lopez-Otin et al., 2023).

Our discovery that mTOR inhibitors are genoprotective make them amenable for use in a wide range of clinical scenarios where the induction of DNA damage leads to pathology. For example, cancer treatments such as radio- or chemotherapy lead to widespread DNA damage of healthy tissue; thus, treatment with a genoprotector, such as low-dose rapamycin, after remission from the original tumour may attenuate the accelerated ageing associated with such cancer therapies (Wang et al., 2024). Likewise, exposure to the space environment, and especially the DNA instability caused by cosmic radiation, is of increasing concern as space travel becomes more commonplace (Beheshti et al., 2021). Our study, through its novel identification of rapamycin as a genoprotector, suggests potential avenues for mitigating these harmful DNA-damaging effects of space travel.

Genoprotectors such as rapamycin present a new and exciting therapeutic approach for the treatment of age-related diseases, both infectious and chronic in nature. SARS-CoV-2, the virus behind the COVID-19 pandemic, induces DNA damage and senescence by degrading DDR enzymes (Gioia et al., 2023); heightened virus-induced senescence in this way strongly contributes to disease mortality (Camell et al., 2021, Lee et al., 2021). Perhaps prophylactic treatment of older care home residents with genoprotectors such as low-dose mTOR inhibitors may provide a much-needed boost to genome stability and immune resilience in this vulnerable population during future pandemics (Cox et al., 2020). Likewise, since mTOR inhibitors improve vaccine responses in older people (Mannick et al., 2018, Mannick et al., 2021, Mannick et al., 2014), future vaccine drives could consider administering short-term mTOR inhibition treatments prior to immunisations against pathogens that particularly affect the older population, such as influenza, coronaviruses, and VZV. Infections by other pathogens, such as *Salmonella* Typhi, *Leishmania*, and some Gram-negative bacteria (which release cytolethal distending toxin), all drive pathology through the induction of DNA damage and senescence (Ibler et al., 2019, Mathiasen et al., 2021, Covre et al., 2018); low-dose mTOR inhibition in these contexts of infection could possibly limit genome instability and disease progression. Though largely untested in humans, research in mice at least suggests that rapamycin improves immune control of Leishmaniasis (Khadir et al., 2018). Finally, in addition to progeroid diseases resulting from a DNA repair deficiency (Werner syndrome, Rothmund-Thomson syndrome, Bloom syndrome, Cockayne’s syndrome, Fanconi’s anaemia, and Ataxia-Telangiectasia), chronic viral infection and rheumatic diseases exhibit decreased DNA repair factor expression in immune cells (Zhao et al., 2018, Shao et al., 2009, Li et al., 2016). It is possible, though unexplored, that rapamycin could limit DNA damage and attenuate pathology in these diseases.

Given the known physiological roles of senescent cells and DNA damage, caution must be taken in administering genoprotective low-dose mTOR inhibitors (de Magalhaes, 2024). For example, inhibiting seno-conversion of virus-infected cells may disrupt their removal by the immune system. Additionally, genoprotectors may disrupt the intentional induction of DNA damage by adaptive immune cells during VDJ recombination, potentially leading to immunodeficiency; however, as the thymus (where T cell VDJ recombination occurs) atrophies with age, it is likely that late-life administration of low-dose rapamycin would not impact on T cells, but possibly B cell maturation. Finally, senescent cells play critical roles in tissue regeneration in response to damage (Chen et al., 2023); thus, genoprotectors may have unforeseen consequences such as by hindering wound healing, a process which is already impaired in older people (Demaria et al., 2014, Wicke et al., 2009).

Taken together, our findings of immune cell benefit on rapamycin treatment *in vitro* and from analysis of PBMCs from an *in vivo* low-dose rapamycin trial, lead us to conclude that rapamycin at 1 mg/day enhances the resilience of the ageing immune system to DNA damage. Our findings support the initiation of phase 2 double-blind placebo-controlled studies of rapamycin to support healthy immunity and reduce immunosenescence in at-risk older adults.

## MATERIALS AND METHODS

### Ethical approval for study

Healthy control blood was taken with fully informed consent under ethical approval from the Local Research Ethics Committee (REC) in Oxford, reference 11/H0711/7, to cover the use of human blood products purchased from National Health Services Blood and Transplant service (NHS England). PBMCs from participants of the Rapamune Trial (registered on Clinical Trials.gov, number NCT05414292) were obtained under ethical approval by the University of Nottingham FMHS REC, reference FMHS 90-0820.

### *In vivo* rapamycin treatment

In the ongoing single-blind, placebo-controlled Rapamune trial (NCT05414292), older male participants (between 50-90 years old) are randomised into two groups and receive either 1 mg/day rapamycin (Pfizer, Belgium) or a placebo sucrose/lactose tablet (Hsconline). The primary outcome is change in muscle mass in response to unilateral leg extension resistance exercise training (RET) 3 times per week for 14 weeks at 75% of the participant’s 1 repetition maximum. Secondary outcomes include changes in muscle strength, power, function, neuromuscular function, and assessment of muscle biopsies on protein synthesis and degradation. In this study, PBMCs were isolated at 5 time points over 4 months of intervention (rapamycin and placebo). After 8 weeks of intervention, the blood concentration of rapamycin and white blood cell counts were measured by Prof. Dan Wilkinson at the University of Nottingham. Inclusion criteria for the trial were that the participant could give informed consent to participate in the study, and that they were able to complete the RET. Exclusion criteria included having a BMI <18 or >35 kg/m^2^, active cardiovascular, cerebrovascular, respiratory, or metabolic disease, clotting dysfunction, having a history of neurological or musculoskeletal conditions, having taken part in a recent study in the last 3 months, or contraindications either to MRI scanning or rapamycin. White blood cell concentrations were quantified in the Royal Derby Hospital Pathology laboratory.

### Measurement of blood rapamycin concentration using LC-MS

50 µl of D3-labelled rapamycin was added to 50 µl of whole blood, before adding 100 µl of precipitation reagent (70:30, Methanol:0.3M Zinc Sulfate). Samples were vortexed for 30 s and mixed on a Vibrax shaker at RT (1000 rpm) for 10 mins. Samples were then centrifuged at 10000g for 10 mins at 4 °C. Supernatant was aliquoted into a 2 ml screw top autosampler vial with low volume insert, before injection into the LC-MS. Samples were quantified against a standard curve of known rapamycin concentrations ranging from 50 ng/ml to 0.39ng/ml prepared in the same way as the samples. Analysis was performed using a Waters ACQUITY UHPLC attached to a Thermo Scientific TSQ Quantum Ultra MS. Rapamycin was isolated using an Agilent Zorbax SB-Aq Narrow Bore RR Column (2.1mm x 100mm x 3.5µm) and a binary buffer system of 2 mM Ammonium Acetate in Water (Buffer A) and 2mM Ammonium Acetate in Methanol (Buffer B) at a flow rate of 0.3 ml/min. Gradient conditions were as follows: 80%B for 0.5 mins, 80%B to 90% B 0.5-2 mins, 99% B 2 – 6 mins, 99%B to 80% 6-6.5 mins, 80% B 6.5-10 mins. Rapamycin was detected using single reaction monitoring (SRM) for m/z transitions of 931.6 m/z – 864.66m/z for unlabelled rapamycin and 934.520 – 864.660 for D3-labelled rapamycin.

### PBMC isolation and culture

Fresh blood was either collected in EDTA tubes (9 ml) or in blood cones (10 ml) as concentrated by-products of the apheresis process, supplied by the National Health Service Blood and Transplant service (NHS England). PBMCs were isolated using standard Ficoll density gradient centrifugation. Briefly, blood was diluted 1:1 in sterile Dulbecco’s PBS (DPBS, 1:5 for blood cones) (Sigma-Aldrich) and 15 ml gently pipetted over 20 ml Histopaque-1077 (Sigma) before centrifugation at 500g for 30 mins at room temperature with minimum deceleration. The PBMC layer was collected by aspiration and washed twice in DPBS. PBMC number was determined by mixing 1:1 (v/v) with Trypan Blue (Sigma) and counting using a haemocytometer. For cryopreservation, PBMCs were resuspended at 5x10^6^ cells/ml in freezing medium (50% FBS, 40% RPMI 1640, 10% DMSO (all Sigma)) and placed at -80°C before transferral to a liquid nitrogen facility (−196°C) for long-term storage. Peripheral blood mononuclear cells (PBMCs) were cultured in R10 (RPMI 1640 (Gibco) containing penicillin (100 U/ml) and streptomycin (100 µg/ml) (both Sigma) and 10% FBS), in a humidified incubator with 5% CO_2_ at 37°C. Details of the drugs used for *in vitro* assays are provided in **Table S1**.

### DNA damage assay in PBMCs

Cryopreserved PBMCs were thawed in R10 (10 ml per 1 cryovial of cells) and centrifuged at 500g for 5 mins. The supernatant was removed, and cells resuspended in R10 at a concentration of 1x10^6^ cells/ml and allowed to recover overnight in R10. The next day, PBMCs were subjected to T cell-specific activation using antibodies targeting CD3 (clone OKT3) and CD28 (clone CD28.2) at 1 µg/ml final concentration (both BioLegend) in the presence of drug treatment or vehicle, where appropriate, for 3 days. Where specified for individual experiments, CD4^+^ T cells were isolated by negative selection on a magnetic column using a kit (Miltenyi) and subsequently activated in wells of a 24-well plate (pre-coated with 1 µg/ml anti-CD3 in PBS for 2 h at 37°C) with 1 µg/ml anti-CD28 in the culture media, for 3 days (0.5-1x10^6^ cells/ml). After 3 days, cells were harvested by trituration, centrifuged at 500g for 5 mins, and resuspended and incubated in R10 with 200 µg/ml zeocin for 2 h or 25 µM H_2_O_2_ for 15 mins (**Table S1**), in a 24-well plate. As both zeocin and H_2_O_2_ are soluble directly in water and cell culture medium, negative controls were not treated with solvent. Cells were then washed in R10 and allowed to recover for a defined period (see individual figures) before being collected for further downstream analysis. Where specified, the DNA damage assay was performed in the presence of rapamycin (10 nM), AZD8055 (100 nM), or chloroquine (10 µM) (drug details provided in **Table S1**).

### Staining for conventional flow cytometry

Cells were pelleted at 500g for 5 mins and resuspended in 50 µl of a master mix of cell-surface-staining antibodies diluted in FACS buffer (0.2% BSA (w/v), 2 mM EDTA in PBS) and Zombie NIR Live/Dead viability dye (1/1000 dilution, BioLegend) with incubation for 30 minutes at 4°C. Cells were washed in FACS buffer and fixed for 20 minutes at 4°C in BD Cytofix Fixation Buffer (BD Biosciences). Permeabilisation of cells was performed by washing cells in 1X BD Phosflow Perm/Wash Buffer I (BD Biosciences), followed by incubation in the permeabilisation buffer for 10 minutes at RT in the dark. Intracellular antibody staining was performed overnight at 4°C in the dark in BD Phosflow Perm/Wash Buffer I. When necessary, staining of unconjugated primary antibodies with a fluorescence-conjugated secondary antibody was performed in BD Phosflow Perm/Wash Buffer I for 1 h at RT in the dark. Cells were washed once more and stored in FACS buffer prior to acquisition on a flow cytometer (BD LSRFortessa) and analysis using FlowJo version 10.9.0 (gating strategy in **Figure S1**). Compensation analysis was performed using single-stained compensation beads (Thermo Fisher or BioLegend). Details of antibodies used for flow cytometry staining are provided in **Table S2**.

### Staining for spectral flow cytometry

PBMCs were stained for analysis for spectral flow cytometry as above, with some modifications. First, cells were harvested and pelleted (500g for 5 mins) and incubated in 50 µl solution containing LIVE/DEAD™ Fixable Blue Dead Cell Stain Kit (1/400 dilution, Invitrogen) and FcR blocking reagent (1/400 dilution, Miltenyi) in PBS for 15 minutes at 4°C. Surface and intracellular antigen staining was then performed as above. After staining, cells were filtered through a 70 µm Flowmi cell strainer (Sigma) to remove aggregates, before analysis on a 5-laser (R/B/V/YG/UV) Aurora spectral flow cytometer (Cytek Biosciences). For each experiment, all samples were processed in one batch and single-colour cell- and bead-based controls were generated in parallel alongside the sample staining for spectral unmixing. Details of antibodies used for spectral flow cytometry staining are provided in **Table S2**. The gating strategy is provided in **Figure S4**.

### Autophagic flux analysis

Measurement of autophagic flux in cells was performed using an antibody-based LC3 assay kit (Cytek Biosciences) with the following modifications (Figure S3). 2 hours prior to LC3 staining, each sample of cells was divided for treatment either with 10 nM Bafilomycin A_1_ (Table S1) or vehicle (0.1% DMSO final concentration) in R10. Cells were harvested and incubated in Live/Dead viability dye (1/1000 dilution, BioLegend) and FcR blocking reagent (1/400 dilution, Miltenyi) in PBS for 15 minutes at 4°C, then surface staining with antibodies was performed as described above, followed by washing in 1X assay buffer (Cytek Biosciences) diluted in dH_2_O, and subsequently permeabilised using 0.05% saponin (w/v) in PBS for 3 minutes at RT. Cells were then incubated with an LC3-FITC conjugated antibody (**Table S2**) in 1X assay buffer for 30 mins at 4°C. For co-staining of LC3 with an anti-γH2AX antibody (**Table S2**), the incubation time was increase to 1 hour in 1X assay buffer. After staining, cells were then washed in 1X assay buffer and fixed in 2% PFA for 10 minutes at RT, before being finally resuspended in FACS buffer and analysed on an LSRFortessa cytometer (BD) or 5-laser Aurora spectral flow cytometer (Cytek) (Figure S3). Autophagic flux was calculated from the mean fluorescence intensity (MFI) of LC3 in Bafilomycin A_1_- and vehicle-treated conditions (*LC3_BafA_* and *LC3_Veh_* respectively) using the formula:

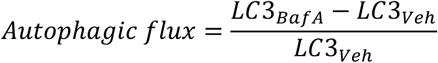

### Cell cycle phase distribution analysis

To assess cell cycle phase distribution using flow cytometry, cells were incubated for 4 h with 10 µM of 5-ethynyl-2′-deoxyuridine (EdU) (Invitrogen). Cells were then harvested and incubated in Live/Dead viability dye (1/1000 dilution, BioLegend) and FcR blocking reagent (1/400 dilution, Miltenyi) in PBS for 15 minutes at 4°C. After this, surface antigen staining for flow cytometry was performed as described above. DNA-incorporated EdU was identified via a click chemistry reaction according to the manufacturer’s instructions (Invitrogen), followed by overnight staining for intracellular γH2AX as described above. Cells were then washed and incubated with FxCycle™ Violet Stain for DNA (1/1000 dilution, Invitrogen) in permeabilisation buffer (from EdU kit, Invitrogen) and analysed on an LSR Fortessa X-20 cytometer (BD).

### Protein synthesis assay

Nascent protein synthesis was measured using the Click-iT™ Plus OPP Alexa Fluor™ 594 Protein Synthesis Assay Kit (Invitrogen). Cells were incubated with 20 µM O-propargyl-puromycin (OPP, from the kit) for 30 mins, then collected and stained for surface antigens. Cells treated for 1 h with 50 µg/ml cycloheximide (**Table S1**) served as a positive control for protein synthesis inhibition. Cells were subsequently fixed, permeabilised, and polypeptide-incorporated OPP was detected, according to the manufacturer’s instructions. Cells were analysed on a Fortessa X-20 flow cytometer (BD).

### Alkaline comet assay

SuperFrost Plus Adhesion slides (VWR) were first pre-coated with normal melting point agarose (NMPA) (1% (w/v) in dH2O, Sigma) and allowed to dry overnight. Following treatments, cells were harvested by trituration and resuspended in PBS at a concentration of 2x10^5^ cells/ml. 250 µl of the cell suspension was mixed with 1 ml of low melting point agarose (1% w/v in PBS, Sigma) pre-warmed to 37°C, and 1 ml of the mixture was pipetted onto one NMPA-coated slide. Coverslips were then placed on the slides and left to gel on ice. For the lysis of cells, coverslips were removed, and slides were incubated overnight at 4°C in fresh, ice-cold lysis buffer (2.5 M NaCl, 100 mM EDTA, 10 mM Tris base containing 1% DMSO (v/v) and 1% Triton X-100 (v/v), pH 10.5). The next day, slides were incubated for 30 mins in the dark with fresh alkaline electrophoresis buffer (300 mM NaOH, 1mM EDTA and 1% DMSO (v/v), pH > 13). Slides were then electrophoresed in the dark for 25 mins at 1 V/cm (distance between electrodes), at a constant current of 300 mA. Slides were then neutralised with 3 x 5 mins incubations in neutralisation buffer (500 mM Tris-HCl, pH 8.1) and subsequently left to dry overnight. The next day, slides were rehydrated for 30 mins in dH_2_O, and DNA was stained for 30 mins with 1X SYBR™ Gold Nucleic Acid Gel Stain (Invitrogen) in dH_2_O. Comets were visualised with a Zeiss Axio Imager and analysed using the OpenComet plugin in Fiji version 2.3.0.

### Statistical tests and figures

All statistical and (log)normality testing of data was performed using GraphPad Prism version 10.0.0. *P*-values indicating statistical significance are either indicated exactly or represented as: ns (not significant) p > 0.05, \**p* ≤ 0.05, \*\**p* ≤ 0.01, \*\*\**p* ≤ 0.001, \*\*\*\**p* ≤ 0.0001. Unless otherwise stated, bar graph data are always represented as mean ± SEM. Box-and-whisker plots always show minimum to maximum values, with the median, 25^th^, and 75^th^ percentiles indicated. Figures were made using Microsoft PowerPoint version 16.88.

## Supporting information

Table S2 - Details of antibodies used in flow cytometry

## ACKNOWLEDGEMENTS

**General:** Investigations into the *in vivo* effects of rapamycin on immune cell senescence were performed on samples from the “Impacts of Mechanistic Target of Rapamycin (mTOR) Inhibition on Aged Human Muscle (Rapamune)” trial, registered on ClinicalTrials.gov, ID NCT05414292: https://clinicaltrials.gov/study/NCT05414292.

## Author contributions

Author contributions were as follows: conceptualization (LK, LSC, AKS, GA, PJA, KS); methodology (LK, EJJ, DJW, NG, GA); investigation (LK, EJJ, DJW, NG); visualization (LK); funding acquisition (LSC, AKS, GA, PJA); project administration (LSC, AKS, GA, PJA, EJJ, LK, DJW, KS); supervision (GA, LSC, AKS); writing – original draft (LK); writing – review & editing (LK, LSC, GA, AKS).

## Data and materials availability

All data are available in the main text or the supplementary materials.

## List of Supplemental Materials

- Figure S1: Gating strategy for CD4^+^ and CD8^+^ T cells using conventional flow cytometry after *in vitro* T-cell-specific activation of PBMCs
- Figure S2: Effects of mTOR inhibitors on human T cell activation over 3 days
- Figure S3: Flow cytometry-based measurement of autophagic flux
- Figure S4: Gating strategy for PBMCs using 27-colour spectral flow cytometry
- Figure S5: *In vivo* rapamycin treatment in older humans
- Table S1: Details of drugs used in cell culture experiments
- Table S2: Details of antibodies used in flow cytometry

