## Supplementary Material for "Rapamycin exerts its geroprotective effects in the ageing human immune system by enhancing resilience against DNA damage"

**CONTENTS**


| Item | Title | Page number |
|---|---|---|
| Figure S1 | Gating strategy for $CD4^+$ and $CD8^+$ T cells using conventional flow cytometry after *in vitro* T-cell-specific activation of PBMCs | 2 |
| Figure S2 | Effects of mTOR inhibitors on human T cell activation over 3 days | 3 |
| Figure S3 | Flow cytometry-based measurement of autophagic flux | 4 |
| Figure S4 | Gating strategy for PBMCs using 27-colour spectral flow cytometry | 5 |
| Figure S5 | *In vivo* rapamycin treatment in older humans | 6 |
| Table S1 | Details of drugs used in cell culture experiments | 7 |
| Table S2 | Antibodies used in study | Separate excel file |

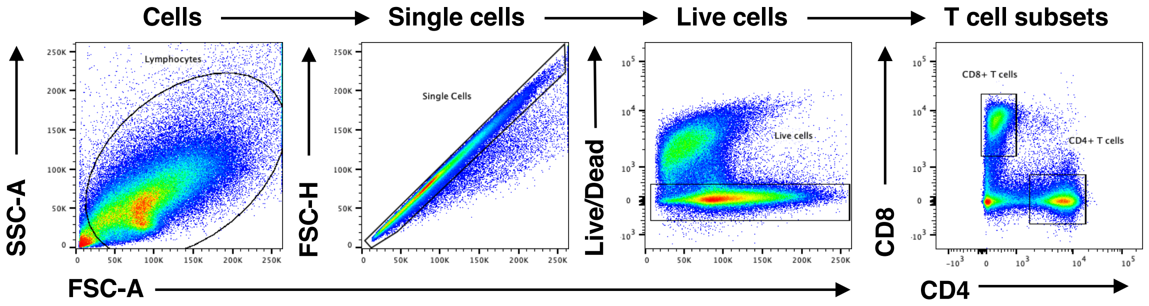


**Figure S1 Gating strategy for CD4+ and CD8+ T cells using conventional flow cytometry after in vitro T-cell-specific activation of PBMCs**

Total cells are gated based on size (forward scatter, FSC-A) and granularity (side scatter, SSC-A) to remove debris. Single cells are determined by their direct proportionality between FSC-area (FSC-A) and FSC-height (FSC-H). Live cells are identified by their absence of fluorescence of a membrane-permeable dye. T cell subsets are then identified by their high expression of either CD4 or CD8.

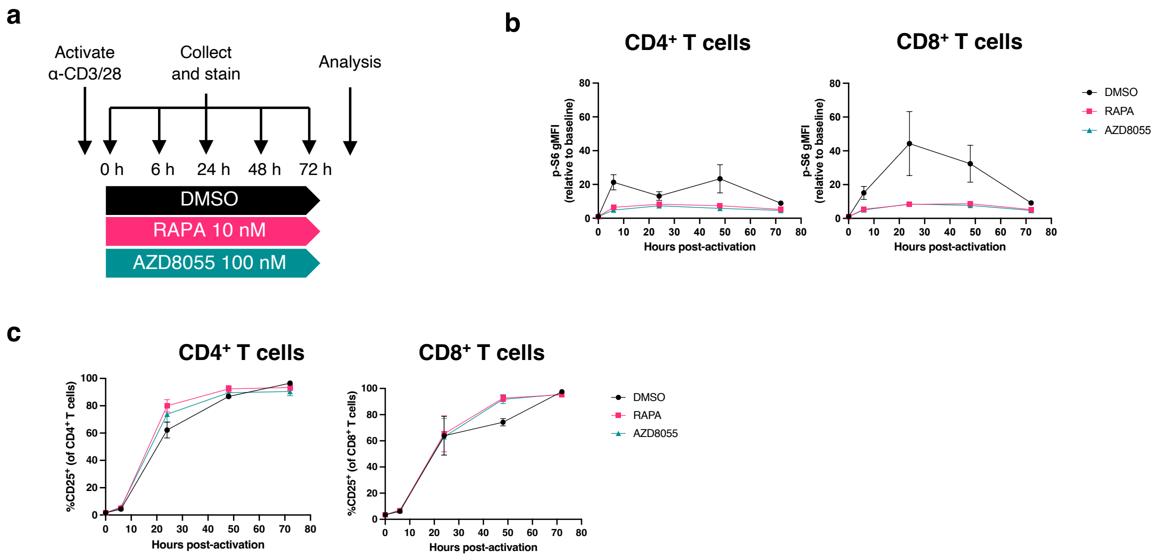


### Figure S2 Effects of mTOR inhibitors on human T cell activation over 3 days

(a) Experimental design for 3-day T-cell-specific activation of PBMCs from healthy donors with 1 µg/ml α-CD3/28 each, in the presence of 10 nM rapamycin (RAPA), 100 nM AZD8055, or DMSO control, with analysis by flow cytometry. (b) p-S6 geometric mean fluorescence intensity (gMFI) in total CD4+ (left) and CD8+ T cells relative to baseline. (c) Proportion of cells positive for (C) CD25 across the 3-day activation in flow cytometry-gated CD4+ or CD8+ T cells.

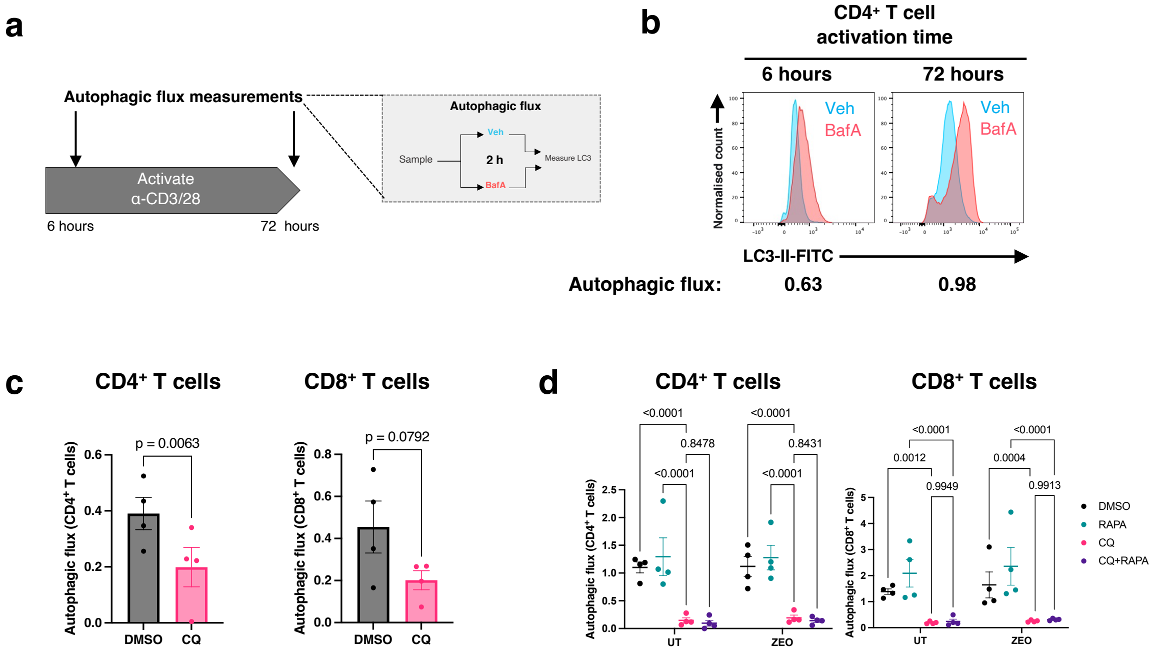


### Figure S3 Flow cytometry-based measurement of autophagic flux

(a) Experimental design in which PBMCs from healthy donors underwent T-cell-specific activation with 1 $\mu$g/ml α-CD3/28. At 6 hours and 72 hours of activation, cells were retrieved and treated with either 10 nM bafilomycin $A_1$ or DMSO vehicle control (Veh) as indicated, and autophagic flux measured using a flow cytometry-based LC3 assay. (b) Representative fluorescence histograms of LC3 levels in gated CD4+ T cells after 6 and 72 hours of activation as in (a), with autophagic flux indicated below. Representative of 3 independent experiments. (c-d) Autophagic flux in CD4+ and CD8+ T cells undergoing 3-day activation in the presence of (c) chloroquine (CQ, 10 $\mu$M) or DMSO vehicle control or (d) following zeocin treatment (or untreated) after 3-day activation in chloroquine (CQ, 10 $\mu$M), rapamycin (RAPA, 10 nM), or both (CQ+RAPA), n=4 healthy donors. P-values are derived from a paired t-test (c) or two-way ANOVA with Tukey's multiple comparisons test (d).

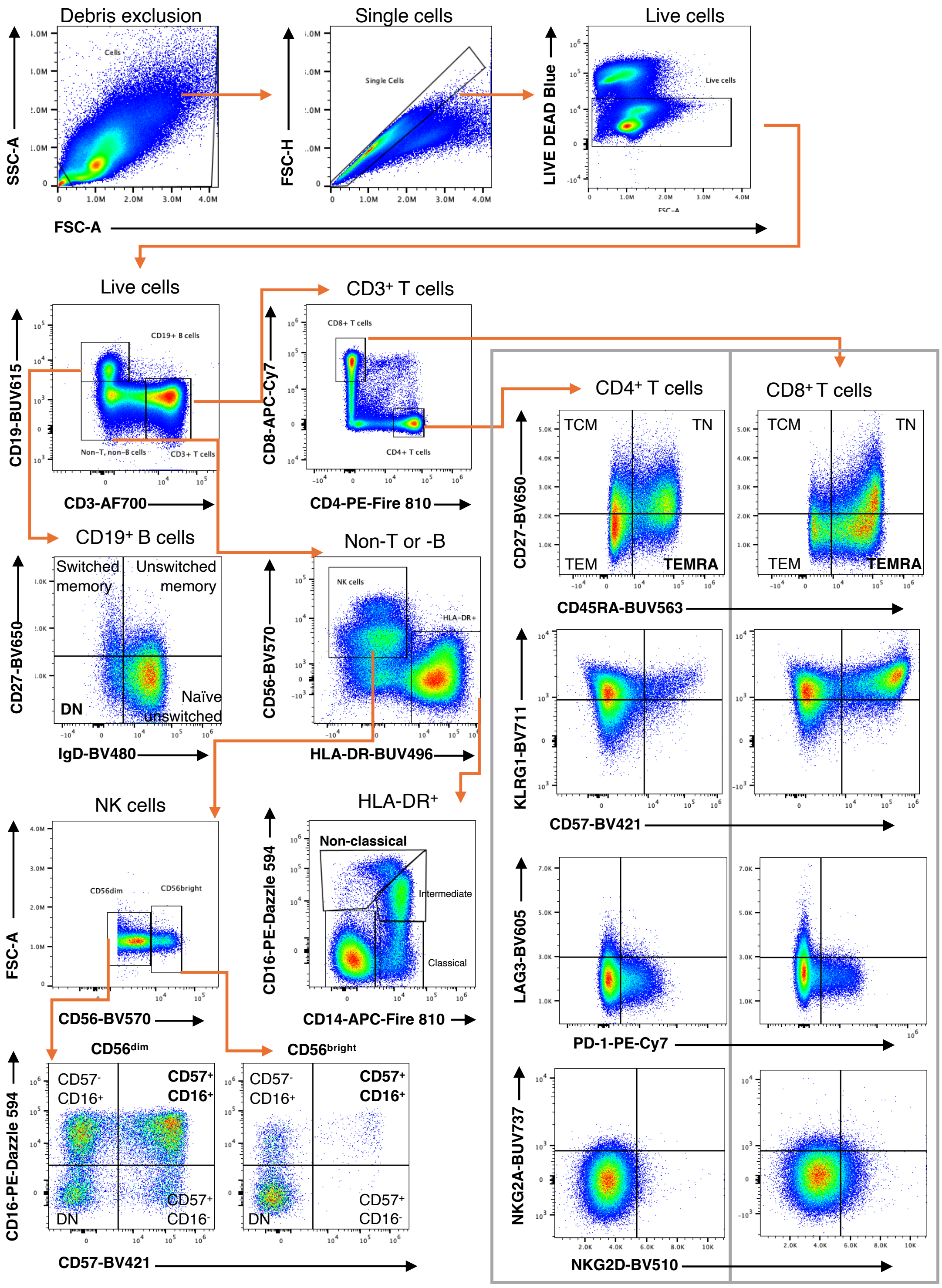


### Figure S4 Gating strategy for PBMCs using 27-colour spectral flow cytometry

Gates are indicated. Orange arrows indicate where a population has been further gated upon.

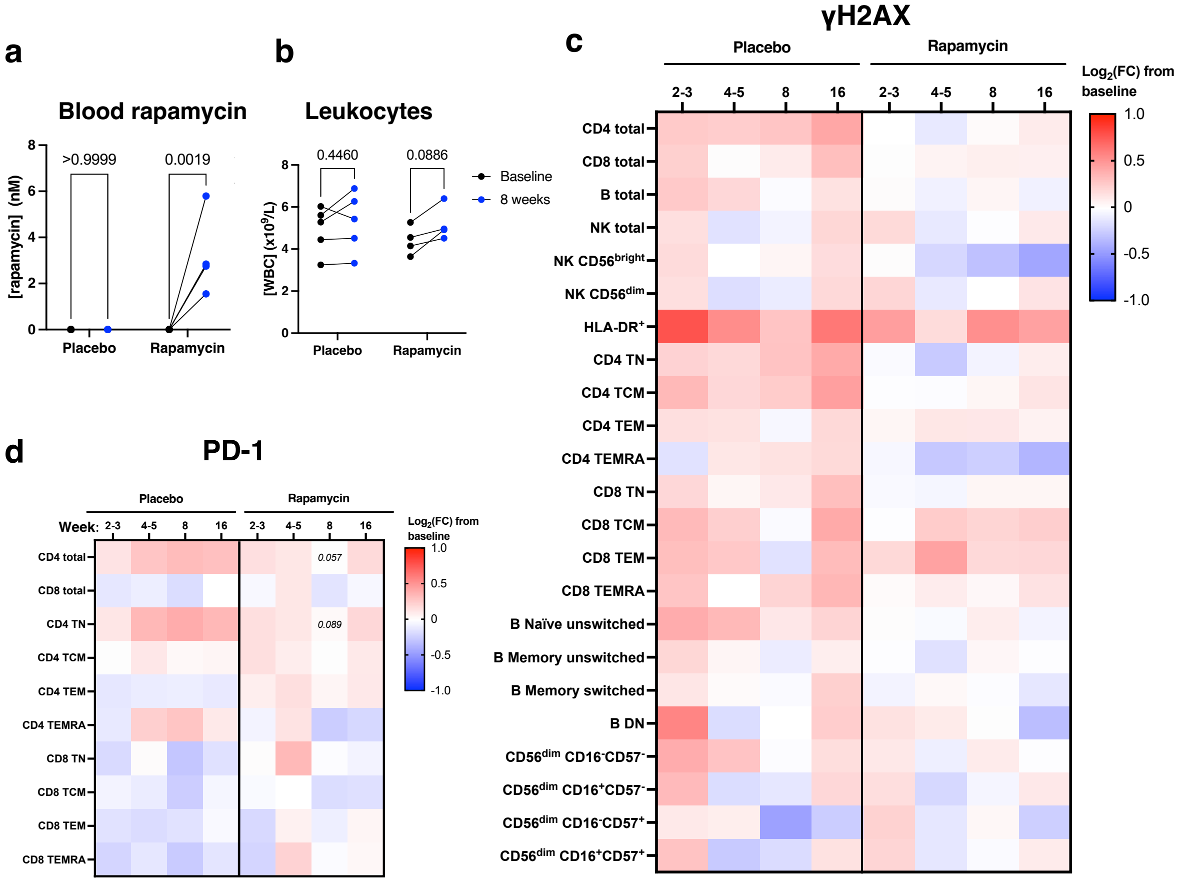


### Figure S5 *In vivo* rapamycin treatment in older humans

(a-b) Blood concentration of rapamycin (a) or white blood cells (b) in participants of placebo and rapamycin groups at week 8 of the study (n=5 placebo, n=4 rapamycin). (c) γH2AX geometric mean fluorescence intensity across immune subsets in the rapamycin trial. (d) Proportion of T cells positive for PD-1 across defined T cell subsets in participants in rapamycin (n=4) and placebo (n=5) groups. In (c-d), each value is expressed as $\log_2$(fold change) from baseline for each participant. Statistical tests in (c-d) are derived from an unpaired t-test between placebo and rapamycin at each time point. *P*-values in (a-b) are determined by two-way ANOVA with Šídák's multiple comparisons tests.

**Table S1 Details of drugs used in cell culture experiments**

| Name | Target | Final concentration | Solvent | Manufacturer | Cat. code |
|---|---|---|---|---|---|
| Zeocin | DSB inducer | 200 $\mu$g/ml | $H_2O$ | Invitrogen | R25001 |
| Rapamycin | mTORC1 inhibitor | 10 nM | DMSO | Alfa Aesar | J67452 |
| AZD8055 | Pan-mTOR inhibitor | 100 nM | DMSO | Stratech Scientific | A8214-APE-10mM |
| Bafilomycin $A_1$ | Autophagy inhibitor | 10 nM | DMSO | Sigma | B1793-10UG |
| Hydrogen peroxide | Oxidative stress | 25 $\mu$M | PBS | Sigma | H1009-100ML |
| Chloroquine | Autophagy inhibitor | 10 $\mu$M | DMSO | Sigma | C6628-25G |
| Cycloheximide | Protein translation inhibitor | 50 $\mu$g/ml | DMSO | Sigma | 239763-M |

N.B. DMSO vehicle controls always contained 0.1% DMSO (v/v) in the cell culture media.

**Table S2 Details of antibodies used in flow cytometry** – see separate excel file.
